# The snoRNP chaperone snR190 and the Npa1 complex form a macromomecular assembly required for 60S ribosomal subunit maturation

**DOI:** 10.1101/2024.09.27.614885

**Authors:** Hussein Hamze, Mariam Jaafar, Ali Khreiss, Carine Dominique, Jessie Bourdeaux, Alfonso Méndez-Godoy, Dieter Kressler, Odile Humbert, Benjamin Albert, Anthony K. Henras, Yves Henry

## Abstract

The early steps of large-ribosomal-subunit assembly feature among the least understood steps of ribosome synthesis in eukaryotes. In *Saccharomyces cerevisiae*, the snR190 box C/D snoRNP chaperone and the Npa1 complex, composed of the α-solenoid scaffold proteins Npa1 and Npa2, the DEAD-box helicase Dbp6, the RNA-binding protein Nop8 and Rsa3, are likely involved in early 25S rRNA folding events. Here, we report for the first time the existence outside pre-ribosomal particles of an independent macromolecular assembly constituted by the Npa1 complex and the snR190 snoRNP chaperone. Nop8 mediates the formation of this assembly and can associate on its own with free snR190. Moreover, Nop8 RRM domain helps tether the snR190 snoRNP to pre-ribosomal particles. snR190 features a specific central stem-loop structure, which is required for high-affinity binding between free snR190 and the Npa1 complex. Deleting this extension does not prevent snR190 association with pre-ribosomal particles but impairs snR190 activity in early pre-rRNA processing events. This work establishes the importance of association with auxiliary protein complexes for optimum snoRNP chaperone activity during rRNA folding events.

## Introduction

The ribosome is the only cellular machine able to catalyse peptide bond formation during protein synthesis. It is a huge ribonucleoprotein particle (RNP) composed of a large and small subunit, each containing ribosomal RNA (rRNA) associated with ribosomal proteins (RPs). The synthesis of ribosomes, to which all cells devote a substantial amount of their resources, involves the production of rRNA precursors (pre-rRNAs), the processing and folding of these precursors and their assembly with RPs to yield mature functional ribosomal subunits. As the ribosome is a ribozyme, a RNA-based enzyme, one key aspect of its synthesis is the acquisition by the rRNAs of their correct three dimensional structure, a prerequisite for catalytic activity. In eukaryotes, ribosome biogenesis starts in the nucleolus with the transcription by RNA polymerase I (RNA Pol I) of a polycistronic precursor to the 18S rRNA (present in the small 40S subunit) and the 5.8S and 25S/28S (yeast/higher eukaryotes) rRNAs (present in the large 60S subunit along with the independently produced 5S rRNA). During transcription, this precursor starts assembling with a subset of RPs as well as scores of so-called assembly factors (AFs) and small nucleolar ribonucleoprotein particles (snoRNPs). These assembly events on the 5’ part of the nascent RNA Pol I transcript, encompassing the 5’ external transcribed spacer and the 18S rRNA sequence (for a cartoon of pre-rRNA sequence organization, see Supplementary Figure S1A), lead to the formation of the first pre-ribosomal particle. This particle, termed in the yeast *Saccharomyces cerevisiae* either 90S or “small subunit processome” (SSU processome) (1,2), can be visualised as terminal knobs in Miller spreads of rDNA chromatin (3). The nascent pre-rRNA may be cleaved co-transcriptionally at site A2 within the internal transcribed spacer 1 (ITS1), which separates the 18S and 5.8S rRNA sequences (for a cartoon of pre-rRNA processing steps in *S. cerevisiae*, see Supplementary Figure S1B). In that scenario, the first independent precursor to the 40S ribosomal subunit, termed the first or primordial pre-40S particle, is released (4). Meanwhile, assembly events proceed on the 3’ part of the nascent pre-rRNA. Once RNA Pol I transcription is completed, this 3’ part encompasses the remaining ITS1, the 5.8S rRNA sequence, the internal transcribed spacer 2 (ITS2), the 25S rRNA sequence and the 3’ external transcribed spacer (3’ETS). The association of AFs, RPs and snoRNPs on this downstream pre-rRNA segment produces the first or primordial pre-60S particle (5). Cleavage of the pre-rRNA transcript may also occur post-transcriptionally. In that case, a huge bi-partite SSU processome/pre-60S particle is generated (5), which will be split into the primordial pre-40S and pre-60S particles by post-transcriptional cleavage within ITS1. The pre-40S and pre-60S particles will then follow independent maturation pathways in the nucleolus, the nucleoplasm and finally in the cytoplasm to yield the mature 40S and 60S ribosomal subunits competent for translation (for a review of these maturation steps, see (6)).

High-resolution cryo-EM structures of many pre-ribosomal particles have been obtained, revealing the folding of rRNAs at different stages of the maturation pathway and the positioning of AFs, allowing predictions to be made as to their modes of action (for a review, see (7)). The entire primordial pre-60S particle has so far escaped high resolution structural characterization, likely due to its high intrinsic flexibility (5). As a consequence, our understanding of the first steps of large-ribosomal-subunit formation lags behind that of other maturation steps. This primordial pre-60S particle contains the 27SA2 pre-rRNA, several box C/D and H/ACA snoRNPs and approximately 40 AFs (5,8). Among these feature four protein modules, namely the Rrp5/Noc1/Noc2 (9), Npa1/Npa2/Nop8/Dbp6/Rsa3 (10,11), Erb1/Ytm1/Nop7 (12,13) and Upa1/Upa2 (5,14) complexes. In addition, no less than seven proteins belonging to related RNA helicase families (Has1, Mak5, Prp43, Dbp3, Dbp6, Dbp7, Dbp9) are present within the primordial pre-60S particle. These could promote pre-rRNA folding (see for example (15)) and/or regulate snoRNA/pre-rRNA interactions. Indeed, we have recently shown that Dbp7 is involved in the removal of the snR190 box C/D snoRNA (16), a prominent component of the primordial pre-60S particle (5). All snoRNPs and many of the AFs present in the primordial particle are absent from its maturation product, the nucleolar pre-60S particle containing the 27SB pre-rRNA. High-resolution cryo-EM data could be obtained for this later particle, which showed that only 5.8S rRNA, ITS2, 5’ domains I and II and part of 3’ domain VI of 25S rRNA were compacted and stabilised at this stage (17–19). Low-resolution cryo-EM data were also collected for the primordial pre-60S particle (5). Comparison with the cryo-EM structure of the later 27SB-containing pre-60S particle (18) allowed the identification of ITS2, 5.8S rRNA and 25S rRNA domains I and II, suggesting that these elements are already stabilised and compacted to a significant extent in the primordial pre-60S particle. This proposal is strengthened by a high-throughput SHAPE probing study (20), which demonstrated that in the primordial pre-60S particle, the ITS2 is already structured, that the root helix of 25S rRNA domain I is formed and that the base-pairing interactions between 5.8S and 25S rRNAs within domain I are already established. Recently, these conclusions were confirmed by a high resolution cryo-EM structure of a nascent primordial pre-60S particle encompassing 5.8S rRNA, ITS2 as well as domains I and II of 25S rRNA (21). Hence, during early steps of large-ribosomal-subunit formation, 25S rRNA domain I is formed and stabilised first, followed by domain II and then VI. These domains form the solvent-exposed face of the large subunit, onto which the other domains can later assemble.

The molecular functions of most AFs present in the primordial pre-60S particle in these early folding steps remain largely elusive. Most are essential for viability and large-ribosomal-subunit synthesis, underscoring their functional importance. At least 14 of them are necessary for normal steady-state accumulation of the 27SA2 pre-rRNA, indicating that they are necessary for the production and/or stability of the primordial pre-60S particle. These include the Rrp5/Noc1/Noc2 complex (22,23), the DExD/H-box proteins Has1 (24), Prp43 (25–27), Dbp3 (28), Dbp7 (29,30), Dbp9 (31), the putative RNA-binding protein Nop4 (32,33) and the Npa1/Npa2/Nop8/Rsa3/Dbp6 (10,11,34,35) complex (in the rest of the manuscript referred to as the “Npa1 complex”). To gain insights into the mechanism of action of these AFs, we and others have searched for their RNA targets by the CRAC UV cross-linking approach (36). Binding sites of Npa1 (10), Prp43 (37) and Rrp5 (38) determined by CRAC have been localised next to the binding site of RP Rpl3 in the root helix of 25S rRNA domain I. Npa1 also binds to 25S rRNA domain VI, adjacent to a second Rpl3-binding site, while Dbp6 and Dbp7 binding sites have been identified in 25S rRNA domains III and V, and V and VI, respectively (15,30). In addition, CRAC data indicate that Npa1, Prp43 and Rrp5 can be cross-linked to some snoRNAs involved in 25S rRNA modification and it has been proposed that Prp43 controls the interaction of a subset of box C/D snoRNAs with 25S rRNA (37). Strikingly, Npa1 interacts predominantly with the box C/D snoRNA snR190, which features two rRNA-complementary sequences allowing it to bind to domain I and to the root helix of domain V. Unlike most snoRNAs, snR190 does not guide rRNA modifications and likely functions as a RNA chaperone. We have shown that lack of snR190 induces a growth defect and impairs the maturation of the first pre-60S particle (16). Altogether, these data suggest that the Npa1 complex collaborates with RNA helicases and the snR190 chaperone in the folding and/or compaction of 25S rRNA domains I, V and VI. The hypothesis of a role for Npa1 in the folding of the 3’ end of 25S rRNA is reinforced by the finding that depletion of URB1, the likely human orthologue of Npa1, altered the pre-rRNA conformation around the 3’end of 28S rRNA (39).

Here, we have investigated the way the Npa1 complex and snR190 interact and the functional implications of this interaction. We provide evidence that snR190 and the Npa1 complex can interact outside pre-ribosomal particles. Nop8 is essential for this interaction. Furthermore, Nop8 or Npa1 depletion prevents snR190 association with pre-ribosomal particles, suggesting that they help tether snR190 to these particles. We also show that a specific stem-loop structure in snR190 is required for high-affinity binding between free snR190 and the Npa1 complex. While this stem-loop structure is not required for snR190 association with pre-60S particles, our results suggest that it is important for snR190 chaperone function during early pre-rRNA processing steps.

## Materials and methods

### Plasmids

Construction of plasmids directing expression of snR190 wild-type and snR190-[mut.AB] mutant was described in (16). Plasmids directing expression of snR190 mutants snR190-[mut.B], snR190-[short Δstem], snR190-[intermediate Δstem], snR190-[large Δstem], were obtained by mutagenesis of the plasmid directing expression of wild-type snR190 (pCH32 vector + wild-type *U14-SNR190* gene insert, (16)) using the PCR-based In-Fusion mutagenesis system (Clontech) and appropriate primers for each mutant (Supplementary Table S1). The original template plasmid was then eliminated by the “Cloning Enhancer” treatment (Clontech) and the linear PCR products were circularized using the In-Fusion system before transformation into competent *E. coli* cells (Stellar, Clontech). To generate a plasmid directing expression of snR190-[mut.B-large Δstem], a mutation in box B was introduced using the above protocol in the plasmid directing expression of snR190-[large Δstem]. All resulting plasmids were verified by sequencing.

### Yeast strains

A BY4742 (*MAT*α, *his3Δ1*, *leu2Δ0*, *lys2Δ0*, *ura3Δ0*) strain expressing Noc1-FPZ (Flag-PreScission cleavage site-tandem IgG-binding Z domains derived from *Staphylococcus aureus* protein A) was produced by transforming BY4742 with a PCR cassette obtained with plasmid pBS1479-NAT-2XFlag-PPX-ZZ and primers listed in Supplementary Table S2. Clones having integrated the nourseothricin resistance gene were selected on YP medium supplemented with 2% glucose and nourseothricin (Jena Bioscience, 80 μg/ml final concentration). *GAL::HA-npa1*/*NOC1::FPZ* and *GAL::HA-nop8*/*NOC1::FPZ* strains (BY4742 background) were produced by transforming the above described Noc1-FPZ-expressing strain with PCR cassettes obtained with plasmid pFA6a-kanMX6-PGAL1-3HA (40) and primers listed in Supplementary Table S2. Clones having integrated the kanMX6 resistance gene were selected on YP medium supplemented with 2% galactose and G418 (Gibco, 200 μg/ml final concentration). A *snR190-[mut.C]* strain (16) expressing Nop8-FPZ was produced as described above using primers listed in Supplementary Table S2. Strains expressing Nop8-HTP (HTP consists of a (His)6 tag, a TEV protease cleavage site and two Z domains from *S. aureus* protein A) or Nop8ΔRRM-HTP under the control of the *NOP8* promoter were obtained upon transformation of the *NOP8* shuffle strain YAM1357 (*MAT***a** *nop8*::HIS3MX4 *ade3*::kanMX4 pHT4467Δ-*NOP8*; W303 background) with plasmid YCplac22-*NOP8*-HTP (pDK10780) or YCplac22-*NOP8.81C*-HTP (pDK10781; 81C denotes that the encoded Nop8ΔRRM starts with amino acid 81 and ends at the native C-terminus) and subsequent counter-selection on plates containing 5-FOA.

Construction of the *RSA3::FPZ*, *DBP6::FPZ, NPA2::FPZ, NOP8::FPZ, GAL::HA-npa1/RSA3::FPZ*, *GAL::HA-npa1/DBP6::FPZ, GAL::HA-npa1/NPA2::FPZ, GAL::HA-npa1/NOP8::FPZ, GAL::HA-nop8/RSA3::FPZ, rrn3.8/RSA3::FPZ* strains was described in (10). Strains *rrn3.8/GAL::HA-npa1/RSA3::FPZ* and *rrn3.8/GAL::HA-npa2/RSA3::FPZ* were produced by transforming the *rrn3.8/RSA3::FPZ* strain with PCR cassettes obtained with plasmid pFA6a-kanMX6-PGAL1-3HA (40) and primers listed in Supplementary Table S2. Strain *rrn3.8/RSA3::FPZ/snR190-[mut.C]* was obtained by crossing haploid strains *rrn3.8/RSA3::FPZ* and *snR190-[mut.C]* (*16*), sporulation of the resulting diploids and tetrad dissections. The correct haploid strains were selected on the basis of temperature sensitivity and resistance to nourseothricin; moreover, mutation of the *SNR190* gene was verified by sequencing a locus-specific PCR product and confirmed by northern analysis. A *snR190-[mut.C]* strain expressing Nop7-TAP (TAP consists of a calmodulin binding peptide tag, a TEV protease cleavage site and two Z domains from *S. aureus* protein A) was constructed by transforming a *snR190-[mut.C]* strain (16) with a PCR fragment obtained using genomic DNA from a *NOP7::TAP* strain (Yeast TAP-tagged ORF collection, Horizon Discovery) and primers listed in Supplementary Table S2.

All strains used in the present work are listed in Supplementary Table S4.

### Immunoprecipitation experiments

Immunoprecipitation experiments with IgG Sepharose were performed as described in (16).

### RNA extractions and northern blot analyses

Yeast total RNAs were extracted as described in (16). Northern blot analyses of high and low molecular weight RNAs were performed as described in (16). Sequences of antisense oligonucleotides used to detect snR190, snR5, snR37 and snR42 snoRNAs are listed in (8,16) and those used to detect snR3 and snR39b snoRNAs in Supplementary Table S3. The sequences of oligonucleotides 23S1 used to detect 35S, 32S and 27SA2 pre-rRNAs and rRNA2.1 used to detect 35S, 32S, 27SA2 and 27SB pre-rRNAs have been reported in (16).

### Tandem affinity purifications

Yeast cell powder was produced using a PM 100 planetary ball mill (Retsch) from a cell pellet obtained from 6 L of yeast culture grown to an OD_600_ of 0.6-0.8. 6 g of cell powder was dissolved in 8 ml of buffer A (20 mM Tris-HCl pH 8, 200 mM KCl, 5 mM MgAc, 0.2% Triton X-100, 1 mM DTT) to which cOmplete EDTA-free protease inhibitor cocktail (Roche) and RiboLock RNase inhibitor (Thermo Scientific) were added. The sample was centrifuged in a Beckman Coulter Optima XE-100 ultracentrifuge at 4°C for 2 h at 39,000 rpm in a Beckman Ti50.2 rotor. The resulting supernatant was subjected to a second ultracentrifugation step during 45 min at 39,000 rpm at 4°C. When tandem affinity purifications were perfomed with *rrn3.8* cells, the two ultracentrifugation steps were omitted and replaced by a centrifugation step in a Beckman Coulter Avanti J-26 XP centrifuge at 4°C for 15 min in a Beckman JA-20 rotor. Then, 9 ml of clarified extract were loaded on a 20 ml column (Bio-Rad) containing 200 μl (bead volume) IgG Sepharose beads (6 Fast Flow, Cytiva). The beads were incubated with the extract during 1 h 30 min at 4°C with gentle agitation. Beads were then washed with 60 ml of buffer B (50 mM Tris-HCl pH 7.4, 200 mM KCl, 5 mM MgAc, 0.2% Triton X-100, 1 mM DTT), then incubated overnight in a 15 ml Falcon tube at 4°C with 50 units of PreScission protease (Cytiva) in 3 ml of buffer B supplemented with 80 units of RiboLock RNase inhibitor (Thermo Scientific). The sample was centrifuged 2 min at 1200 rpm in an Eppendorf 5810 R benchtop centrifuge and the supernatant was collected. The IgG Sepharose beads were mixed with 2 ml of buffer B and centrifuged 2 min at 1200 rpm in an Eppendorf 5810 R benchtop centrifuge. The supernatant was collected and mixed with the previously collected supernatant. The resulting sample was applied to a 10 ml Bio-Rad column containing 100 μl (bead volume) of anti-Flag M2 affinity gel (Sigma) and incubated with the gel for 1 h at 4°C on a rotating wheel. The column was washed with 40 ml of buffer C (10 mM Tris-HCl pH 7.4, 150 mM NaCl, 5 mM MgCl2, 1 mM DTT). Purified complexes were eluted by adding five times 200 μl of buffer C supplemented with 2x Flag peptide (400 μg/ml, IGBMC Strasbourg). Alternatively, the gel was mixed with 1 ml of buffer C and two 550 μl aliquots were transferred to two 1.5 ml Eppendorf tubes. The tubes were centrifuged 1 min, 2000 rpm in an Eppendorf 5415 R benchtop centrifuge and the supernatant was discarded. RNAs were extracted from the bead pellet of one sample. The beads of the other sample were directly mixed with 50 μl of SDS sample buffer (100 mM Tris-HCl, pH 8.0, 4% SDS, 20% glycerol, 0.04% bromophenol blue and 200 mM DTT) for protein analysis.

### pCp labelling

RNAs were labelled with 10 μCi [^32^P]pCp (PerkinElmer) and 10 units of T4 RNA ligase (Thermo Scientific) in the presence of 40 units of RNasin ribonuclease inhibitor (Promega) in a 20 μl reaction volume at 4°C overnight. Labelled RNAs were phenol/chloroform extracted, ethanol precipitated, washed and resuspended in 50% formamide (v/v). The labelled RNAs were electrophoresed in Tris-Borate EDTA buffer on a 6% acrylamide/bis-acrylamide (19:1)/50% urea (w/v) gel for 3 h at 1800 V. The gel was dried on 3MM paper and exposed to a phosphorimager screen.

### Western blot analyses

Western blot analyses were performed as described in (15,16). FPZ-tagged or TAP-tagged proteins were detected with rabbit peroxidase anti-peroxidase (PAP) soluble complex (Dako) (1:10,000 dilution). Primary antibodies used to detect Npa1 (1:2000 dilution), Dbp6 (1:10,000 dilution), Nop8 (1:1000 dilution), Nhp2 (1:5000 dilution) were generated in rabbits by custom antibody production services and described elsewhere (10,41). Yeast Nop1 was detected with anti *Xenopus laevis* Fibrillarin antibodies (1:270 dilution) raised in rabbits. Anti-rabbit IgG-HRP conjugate (Promega) were used when needed as secondary antibodies (1:10,000 dilution).

## Results

### Npa1 or Nop8 depletion perturbs the association of snR190 with pre-ribosomal particles

We previously showed that snR190 inactivation affects the association of Npa1 complex members with pre-ribosomal particles (16). We wanted to determine whether the reverse was also true, i.e., whether Npa1 complex inactivation impacts snR190 association with pre-ribosomal particles. To test this hypothesis, we immunoprecipitated 90S and early pre-60S pre-ribosomal particles using tagged Noc1 (Noc1-FPZ) as bait and extracts from wild-type cells and cells depleted of Npa1, which is essential for Npa1 complex formation (10). The co-precipitation efficiency of snR190 was then analysed by northern (Figure 1A). As controls, we also assessed the co-precipitation efficiency of a subset of snoRNAs, which are either efficiently cross-linked to Npa1 (snR5 and snR42), moderately so (snR39b) or hardly at all (snR3) in CRAC experiments (10). Npa1 depletion decreased the co-precipitation efficiency of all snoRNAs tested (Figure 1A). This was neither due to reduced precipitation of tagged Noc1 (Figure 1A), nor to lack of co-precipitation of pre-rRNAs (Supplementary Figure S2A) in absence of Npa1. Strikingly, the co-precipitation efficiency of snR190 was reduced the most in absence of Npa1 (10-fold reduction), while that of the other snoRNAs was significantly less affected (2- to 3-fold reduction) (Figure 1A).

**Figure 1.**
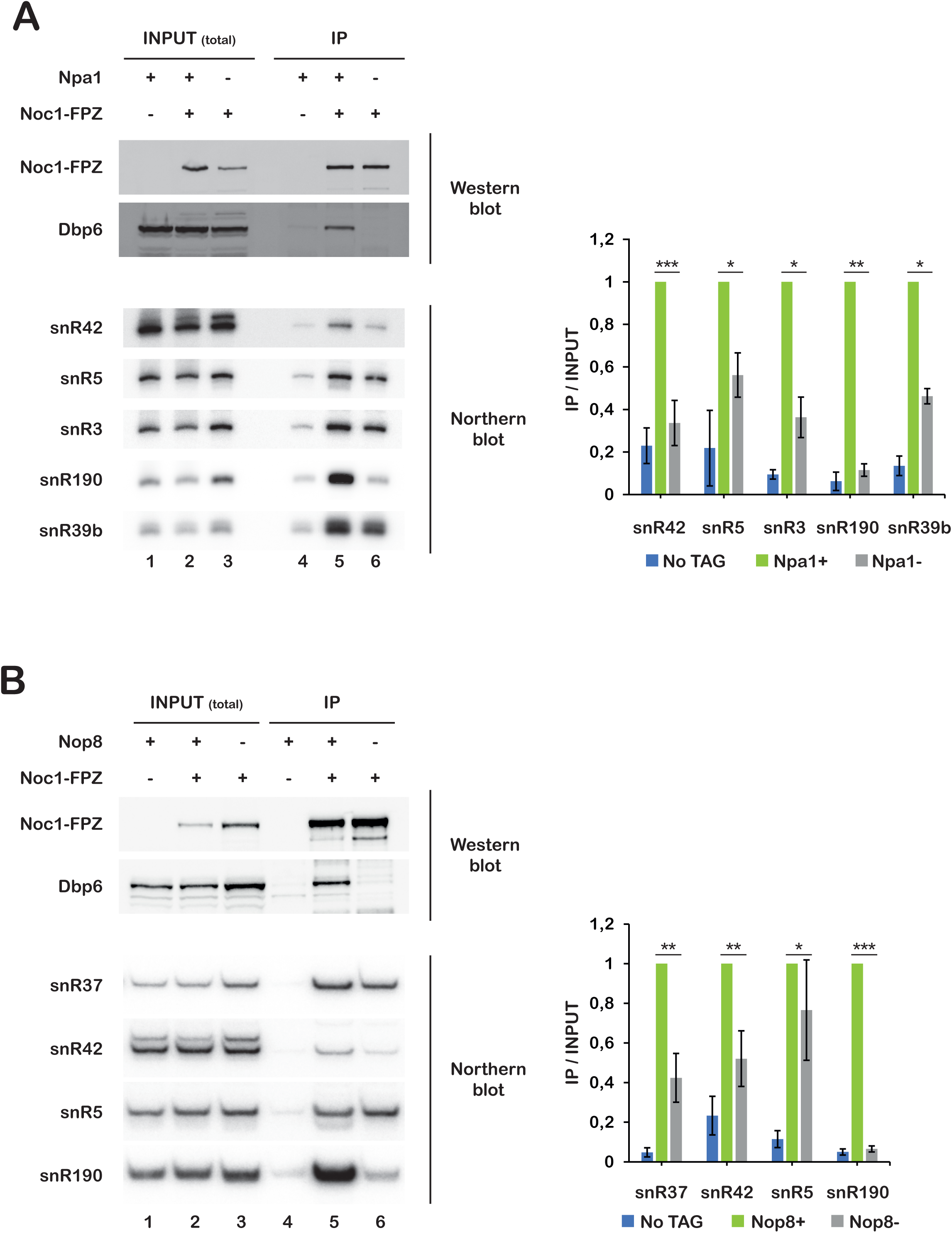
Npa1 or Nop8 depletion strongly affects the association of snR190 with pre-ribosomal particles. Immunoprecipitation experiments were carried out using IgG Sepharose and extracts from a BY4742 wild-type strain (labelled Npa1 +, Noc1-FPZ - (**A**) or Nop8 +, Noc1-FPZ - (**B**)), a strain expressing Noc1-FPZ (i.e., Noc1 bearing a C-terminal Flag tag, followed by a PreScission cleavage site and two IgG-binding Z domains of *S. aureus* protein A) (labelled Npa1 +, Noc1-FPZ + (**A**) or Nop8 +, Noc1-FPZ + (**B**)), a *GAL::HA-npa1*/*NOC1::FPZ* strain (labelled Npa1 -, Noc1-FPZ + (**A**)) or a *GAL::HA-nop8*/*NOC1::FPZ* strain (labelled Nop8 -, Noc1-FPZ + **(B**)) expressing Noc1-FPZ and depleted of either Npa1 or Nop8, respectively, following growth in glucose-containing medium for 14 hours. Total proteins or RNAs were extracted from input extracts (INPUT (total)) or from immunoprecipitated samples (IP) and analysed by western (top panels) or northern (bottom panels), respectively. Noc1-FPZ was detected using PAP, Dbp6 with specific antibodies. The indicated snoRNAs were detected by northern using antisense oligonucleotide probes. Quantifications of northern data are presented in the histograms on the right. Ratios of precipitated snoRNAs versus total snoRNAs present in the input extracts (IP/INPUT) were computed from phosphorimager scans of northern membranes. Ratios obtained for the non-depleted strains expressing Noc1-FPZ were arbitrarily set at 1. Error bars correspond to standard deviations computed from three independent biological replicates. Statistically significant differences determined using one-tailed Student’s t-test are indicated by asterisks (***= p<0.001; **= p<0.01; *= p<0.1).

We showed previously that in contrast to Rsa3, Npa2 and Dbp6, the association of Nop8 with pre-ribosomal particles is not strongly affected by Npa1 depletion (10). We therefore assessed whether lack of Nop8 also has a specific impact on snR190 association with pre-ribosomal particles. Strikingly, while Nop8 depletion did not affect precipitation of Noc1-FPZ (Figure 1B) nor the co-precipitation of pre-rRNAs (Supplementary Figure S2B), it reduced snR190 co-precipitation to background levels (Figure 1B). The co-precipitation of other snoRNAs tested was also affected, but to a far lesser extent.

We also analysed the effect of depleting Dbp6, the DEAD-box ATPase component of the Npa1 complex (15), on snR190 association with pre-ribosomal particles. Contrary to the lack of Nop8, Dbp6 depletion had little effect on snR190 co-precipitation with Noc1-FPZ (Supplementary Figure S3).

We conclude that Npa1 and Nop8 are specifically required for the stable association of snR190 with pre-ribosomal particles, while the enzymatic activity of Dbp6 is not. Besides Npa1, for which a direct interaction with snR190 could be revealed by CRAC (10), Nop8 therefore also seems to play a key role in snR190 integration and/or retention within early pre-60S pre-ribosomal particles.

### snR190 interacts with the Npa1 complex outside pre-ribosomal particles

The mutual dependence of the Npa1 complex and snR190 for their stable association with pre-ribosomal particles ((16) and this work) may be due to the fact that they are integrated as a pre-formed module into the particles. We therefore tested whether snR190 is associated with the Npa1 complex outside pre-ribosomal particles. To purify the free Npa1 complex, we used a strain in which RNA Pol I can be inactivated conditionally, abrogating *de novo* pre-ribosomal particle production. This strain (*rrn3.8*) expresses a temperature-sensitive version of the Rrn3 RNA Pol I transcription factor. At 25°C, RNA Pol I is active and pre-ribosomal particles are produced and matured. After transfer for four hours to 37°C, RNA Pol I transcription is shut down and pre-ribosomal particles are no longer detectable (Supplementary Figure S4A). We proceeded to tandem affinity purification of the Npa1 complex using an FPZ-tagged version of Rsa3 from *rrn3.8* cells grown at 37°C. Western analysis indicated that the integrity of the Npa1 complex was maintained in this condition (Supplementary Figure S4B) and strikingly, only snR190 was co-purified, as shown by northern (Figure 2A) and pCp labelling (Figure 2B). These data indicate that snR190 can specifically interact with the Npa1 complex when it is not associated with pre-ribosomal particles. To rule out the possibility that such interaction only occurs in cells in which RNA Pol I transcription has been abolished, we repeated the tandem affinity purification of the Rsa3-FPZ bait protein from an extract of a wild-type strain that was subjected to two consecutive ultracentrifugation steps to pellet pre-ribosomal particles, as previously described (10) (see also Supplementary Figure S5A). Northern analysis showed that only snR190 was specifically co-purified with tagged Rsa3 under these conditions (Supplementary Figures S5B and C).

**Figure 2.**
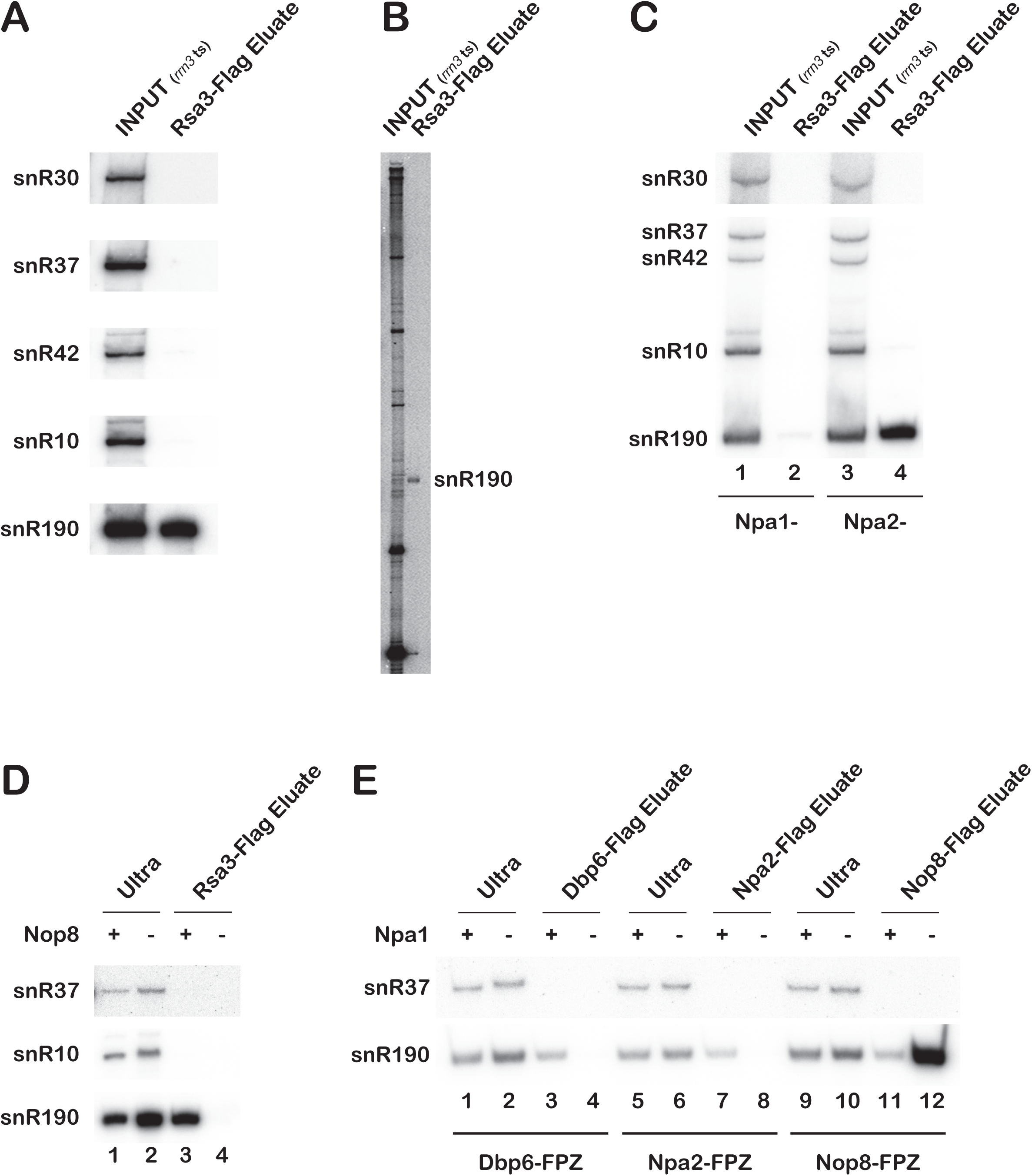
(**A-B**) snR190 specifically interacts with the isolated Npa1 complex. Rsa3-FPZ was subjected to tandem affinity purification from extracts of *rrn3.8/RSA3::FPZ* cells grown at 37°C for 4 hours. RNAs from the initial total clarified extract (INPUT (*rrn3* ts)) or from the eluate from the anti-Flag affinity column (Rsa3-Flag Eluate) were analysed by northern (**A**) or pCp labelling (**B**). (**C**) Nap1, but not Npa2, is required for the interaction between tagged Rsa3 and snR190. Rsa3-FPZ was subjected to tandem affinity purification from extracts of *rrn3.8/GAL::HA-npa1/RSA3::FPZ* (Npa1 -, lanes 1 and 2) or *rrn3.8/GAL::HA-npa2/RSA3::FPZ* (Npa2 -, lanes 3 and 4) strains grown in glucose-containing medium at 37°C for 8 hours. RNAs from the initial total clarified extracts (INPUT (*rrn3* ts)) or from the eluates from the anti-Flag affinity columns (Rsa3-Flag Eluate) were analysed by northern with probes complementary to the indicated snoRNAs. (**D**) Nop8 is required for the interaction between tagged Rsa3 and snR190. Tandem affinity purification of Rsa3-FPZ was carried out with soluble extracts, obtained after two ultracentrifugation steps, from *RSA3::FPZ* (Nop8 +) or *GAL::HA-nop8/RSA3::FPZ* (Nop8 -) cells grown in glucose-containing medium for 14 hours. RNAs from these extracts (Ultra) or from the anti-Flag affinity column eluate (Rsa3-Flag Eluate) were analysed by northern with probes complementary to the indicated snoRNAs. (**E**) Nop8 interacts with snR190 in absence of Npa1 complex formation. Tandem affinity purification of FPZ-tagged proteins was carried out with soluble extracts, obtained after two ultracentrifugation steps, from *DBP6::FPZ, NPA2::FPZ, NOP8::FPZ* cells (all labelled ‘Npa1 +’) or *GAL::HA-npa1/DBP6::FPZ, GAL::HA-npa1/NPA2::FPZ, GAL::HA-npa1/NOP8::FPZ* cells (all labelled ‘Npa1 -’) grown in glucose-containing medium for 14 hours. RNAs from these extracts (Ultra) or from the anti-Flag affinity column eluate (Flag Eluate) were analysed by northern with probes complementary to the indicated snoRNAs.

Given the specific association between snR190 and the isolated Npa1 complex, we assessed whether snR190 contributes to the cohesion of the free Npa1 complex. We performed a tandem affinity purification of tagged Rsa3 from *rrn3.8* cells expressing or lacking snR190 and grown at 37°C. The co-purification of Npa1 and Dbp6 with tagged Rsa3 is not diminished by the absence of snR190 (Supplementary Figure S6A). Strikingly, as previously observed (16), lack of snR190 induces a clear drop in the steady-state accumulation of Nop8 (Supplementary Figure S6A, compare lanes 1 and 2). Nevertheless, Nop8 is co-purified with tagged Rsa3 in absence of snR190. These data suggest that snR190 is not required for the association between Rsa3 and at least Npa1 and Dbp6 outside pre-ribosomal particles.

The co-purification with Rsa3 of the methyltransferase Nop1, a known component of the snR190 snoRNP (42), only in the presence of snR190, but not of Nhp2, a core component of box H/ACA snoRNPs (43), strengthens the conclusion that the snR190 snoRNP associates with the free Npa1 complex (Supplementary Figure S6B).

### The snR190 snoRNP is tethered to the free Npa1 complex via Npa1 and Nop8

CRAC data indicated that Npa1 directly binds to snR190 (10), strongly suggesting that the snR190 snoRNP is tethered to the Npa1 complex via at least Npa1. We next assessed the contribution of the other members of the Npa1 complex to the interaction with snR190. We first analysed the consequence of Npa2 depletion, or Npa1 as control, on the ability of snR190 to interact with Rsa3-FPZ when *de novo* ribosome biogenesis is inhibited. For this analysis, we used *rrn3.8/GAL::HA-npa1/RSA3::FPZ* or *rrn3.8/GAL::HA-npa2/RSA3::FPZ* strains grown in glucose-containing medium at 37°C to inactivate RNA Pol I transcription and deplete Npa1 or Npa2, respectively. When Npa1 was depleted, Npa1 complex formation was inhibited, as also shown previously (10) and snR190 failed to be co-purified with Rsa3 (Supplementary Figure S7A, Figure 2C, lanes 1 and 2). In absence of Npa2, an Npa1/Nop8/Rsa3 sub-complex could form, consistent with our previous findings (10) and snR190 was efficiently co-purified with this module (Supplementary Figure S7B; Figure 2C, lanes 3 and 4). Thus, Npa2 is dispensable for snR190 interaction with the Npa1 complex.

Given the importance of Nop8 for snR190 association with pre-ribosomal particles (see Figure 1B), we next assessed the effect of Nop8 depletion on the ability of snR190 to interact with the isolated Npa1 complex. Tandem affinity purification of Rsa3-FPZ from Nop8-depleted cells after two ultracentrifugation steps to pellet pre-ribosomal particles showed that snR190 does not co-purify with the Npa1/Npa2/Dbp6/Rsa3 sub-complex (Supplementary Figure S7C, Figure 2D). These results indicate that Nop8 is essential for the interaction of snR190 with the free Npa1 complex.

The above data point out that Rsa3 on its own cannot interact with snR190. To determine whether the other Npa1 complex members, Dbp6, Npa2 or Nop8 can interact on their own with the free snR190 snoRNP, we depleted Npa1 to prevent Npa1 complex formation and assessed whether snR190 could be co-purified with FPZ-tagged Dbp6, Npa2 or Nop8 from soluble extracts obtained after two ultracentrifugation steps to pellet pre-ribosomes (Supplementary Figure S7D). Northern data indicated that when Npa1 is expressed, snR190 was specifically co-purified with tagged Dbp6, Npa2 and Nop8 (Figure 2E, lanes 3, 7 and 11), further strengthening the conclusion that snR190 is specifically associated with the isolated Npa1 complex. In contrast, Npa1 depletion severely reduced the efficiency of snR190 co-purification with tagged Dbp6 or Npa2 (Figure 2E, lanes 4 and 8). Strikingly however, the interaction between tagged Nop8 and snR190 was enhanced in absence of Npa1 (Figure 2E, compare lanes 11 and 12), strongly suggesting that Nop8 directly binds to the snR190 snoRNP.

We conclude from these analyses that the snR190 snoRNP is tethered to the free Npa1 complex via Npa1 itself and Nop8.

### Nop8 RNA Recognition Motif (RRM) strengthens snR190 association with pre-ribosomal particles

Nop8 contains a RNA recognition motif (RRM) at its N-terminus (44,45). We reasoned that Nop8 might connect the Npa1 complex to snR190 via this RRM. To test this hypothesis, we performed immunoprecipitation experiments with total extracts of cells expressing tagged versions of Nop8 lacking the RRM (Nop8ΔRRM), or wild-type Nop8 as control and assessed the extent of snR190 co-precipitation. Nop8ΔRRM accumulated and was precipitated to the same extent as full-length Nop8 (Figure 3A) and snR190 was co-precipitated with Nop8ΔRRM nearly as well as with full-length Nop8 (Figure 3B). In contrast, the co-precipitation efficiency of snR37 with Nop8ΔRRM dropped 3-fold compared with the wild-type situation, which might reflect a decreased co-precipitation of pre-ribosomal particles. Indeed, high molecular weight northern data showed that deletion of the RRM reduced the co-precipitation efficiency of the 35S, 32S and 27SA2 pre-rRNAs approximately 2-fold (Figure 3C). These data demonstrate that the RRM motif is dispensable for Nop8 interaction with snR190 but likely strengthens its association with the pre-rRNAs. Given that Nop8 is crucial for snR190 association with pre-ribosomal particles, we next tested whether the deletion of its RRM reduces snR190 pre-ribosome association. We immunoprecipitated pre-ribosomal particles via Noc1-FPZ from extracts of *GAL::HA-nop8/NOC1::FPZ* cells depleted of genome-encoded Nop8 and expressing Nop8ΔRRM, or, as control, wild-type Nop8 from plasmids. Nop8ΔRRM displayed reduced co-precipitation with Noc1-FPZ (Supplementary Figure S8A), suggesting weakened association with pre-ribosomal particles. Co-precipitation of snR190 with Noc1-FPZ dropped 3-fold under conditions of Nop8ΔRRM expression, providing evidence of weakened snR190 association with pre-ribosomal particles (Supplementary Figure S8B). Interestingly, the weakened association of Nop8ΔRRM, and as a consequence of snR190, with pre-ribosomal particles was correlated with a pre-rRNA processing phenotype, characterized by a 30% drop of the 27SB pre-rRNA/27SA2 pre-rRNA ratio when compared to the wild-type control (Supplementary Figure S8C).

**Figure 3.**
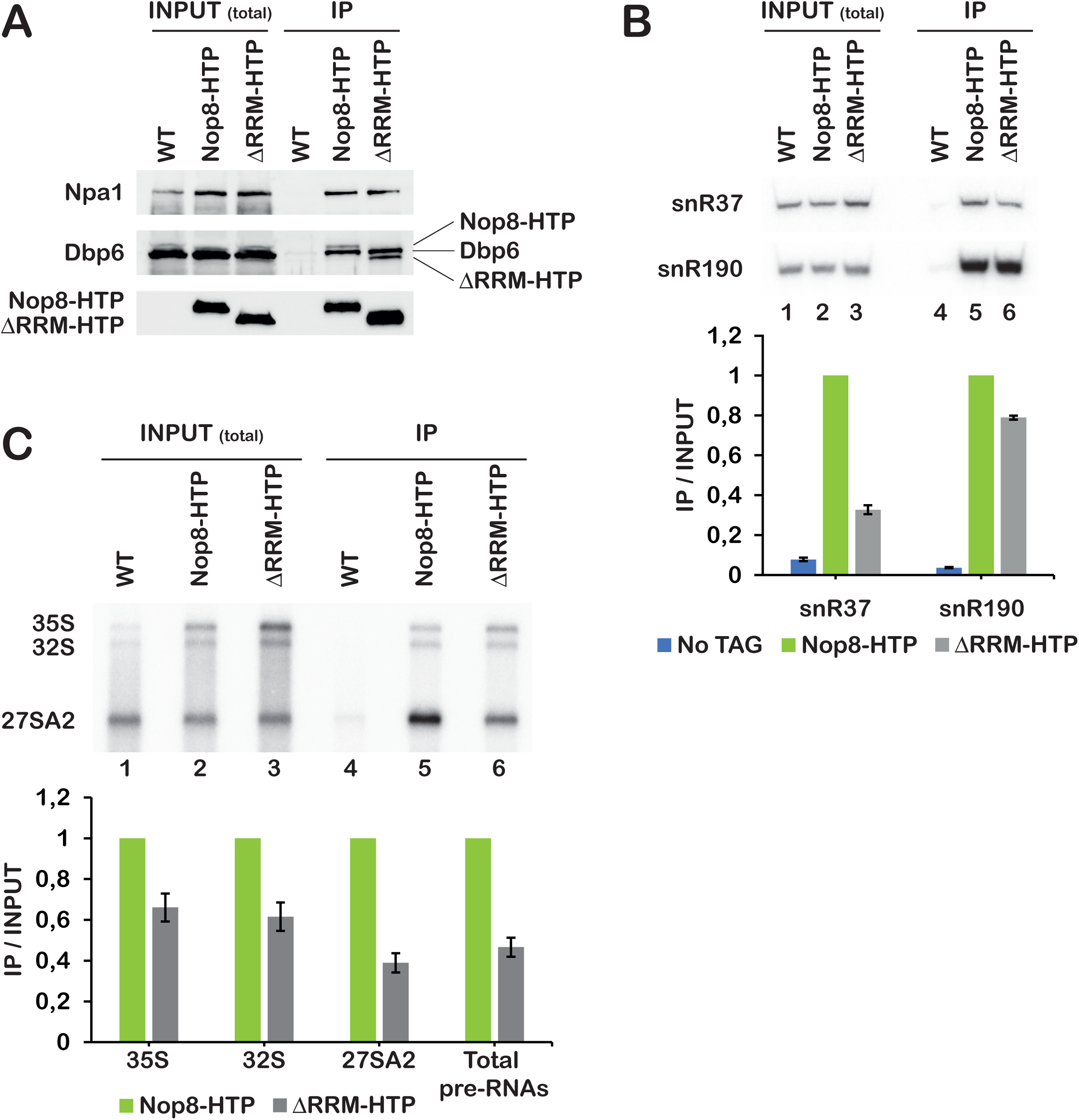
The RRM domain is dispensable for Nop8 interaction with snR190 but strengthens Nop8 association with pre-rRNAs. Immunoprecipitation experiments were carried out using IgG Sepharose and extracts from the parental W303 wild-type strain (WT), a strain expressing Nop8-HTP (i.e., Nop8 bearing a C-terminal (His)6 tag, followed by a TEV cleavage site and two IgG-binding Z domains of *S. aureus* protein A) or Nop8ΔRRM-HTP (ΔRRM-HTP). Total proteins or RNAs were extracted from input extracts (INPUT (total)) or from immunoprecipitated samples (IP) and analysed by western (**A**) or northern (**B, C**). (**A**) Nop8-HTP and Nop8ΔRRM-HTP were detected using PAP, Npa1 and Dbp6 with specific antibodies. (**B**) The indicated snoRNAs were detected with specific antisense oligonucleotide probes. Quantification of northern data is presented in the histogram below the northern. Ratios of precipitated snoRNAs versus snoRNAs present in the input extracts (IP/INPUT) were computed from phosphorimager scans of northern membranes. Ratios obtained for the strain expressing Nop8-HTP were arbitrarily set at 1. Error bars correspond to standard deviations computed from two technical replicates. (**C**) The indicated pre-rRNAs were detected using the 23S1 probe. Quantification of northern data is presented in the histogram below the northern. Ratios of precipitated pre-rRNAs versus input pre-rRNAs (IP/INPUT) were computed from phosphorimager scans of northern membranes. Ratios obtained for the strain expressing Nop8-HTP were arbitrarily set at 1. Error bars correspond to standard deviations computed from two technical replicates.

### The internal stem-loop structure of snR190 plays a key role in its interaction with the isolated Npa1 complex

We next investigated which sequences of snR190 are necessary for its specific interaction with the isolated Npa1 complex. According to our previous Npa1 CRAC data set (10), Npa1 cross-linking sites are positioned within an internal stem-loop extension and within box B, which is complementary to 25S rRNA domain I (Supplementary Figure S9A). We therefore analysed the contribution of these regions of snR190 to its interaction with the isolated Npa1 complex. We generated plasmids directing expression of snR190 mutants featuring nucleotide substitutions at the Npa1 cross-linking sites within box B (snR190-[mut.B]), progressive truncations of the internal stem-loop (snR190-[short Δstem], snR190-[intermediate Δstem], snR190-[large Δstem]), or featuring the box B mutations and lacking the internal stem-loop (snR190-[mut.B-large Δstem]). snR190-[short Δstem] retains all Npa1 cross-linking sites, while one cross-linking region is absent in snR190-[intermediate Δstem] and two in snR190-[large Δstem]. We also included in our analysis a mutant of snR190 featuring mutations within boxes A and B (snR190-[mut.AB]) that abolish their complementarity to 25S rRNA domains I and V (16). These snR190 mutations do not reduce snR190 steady-state levels (Supplementary Figure S9B). We then assessed the ability of snR190 mutants to interact with the isolated Npa1 complex. To do so, plasmids directing expression of wild-type snR190 or the above-described mutants were transformed into the *rrn3.8/RSA3::FPZ/snr190-[mut.C]* strain (which does not express snR190 due to mutation of conserved box C, (16)). Rsa3-FPZ was precipitated from extracts of the transformed strains grown at 37°C to inactivate *de novo* ribosome biogenesis. Strikingly, the co-precipitation efficiency of snR190 mutants lacking the internal stem-loop (snR190-[large Δstem] and snR190-[mut.B-large Δstem]) was reduced 10-fold relative to the wild-type situation (Figure 4A, lanes 4 and 8). The co-precipitation efficiency of snR190-[intermediate Δstem] was also reduced, but to a far lesser extent. We conclude that the isolated Npa1 complex interacts with snR190 via its internal stem-loop structure.

**Figure 4.**
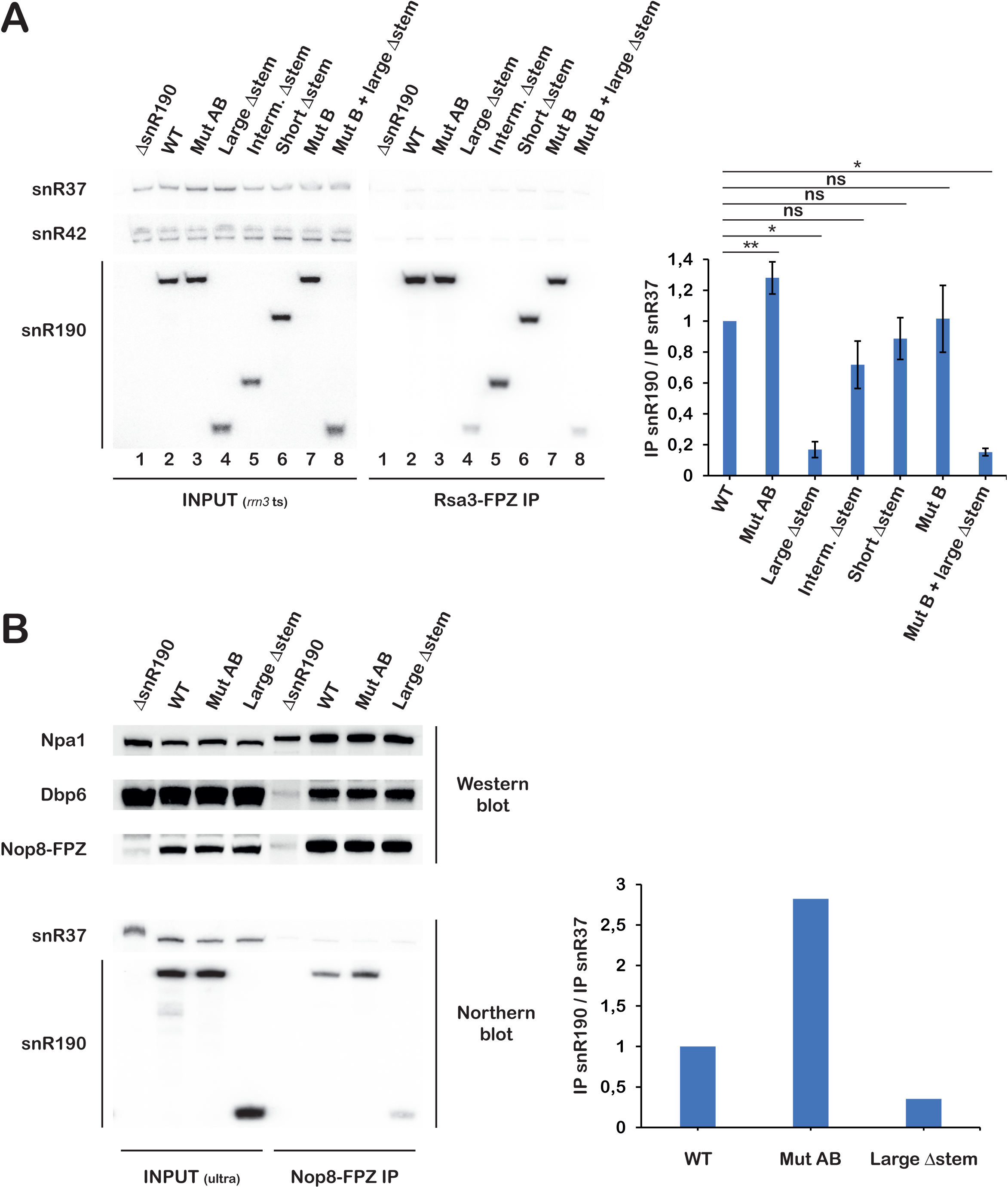
The internal stem-loop of snR190 is crucial for the interaction of snR190 with the isolated Npa1 complex. (**A**) The *rrn3.8/RSA3::FPZ/snr190-[mut.C]* strains transformed with plasmids directing expression of wild-type snR190 (WT), the indicated snR190 mutants or the empty vector (ΔsnR190) were grown for 4 hours at 37°C to inactivate RNA Pol I transcription. Immunoprecipitation experiments were carried out with IgG Sepharose to precipitate Rsa3-FPZ. RNAs extracted from the total cellular extracts (INPUT (*rrn3* ts)) or immunoprecipitated samples (Rsa3-FPZ IP) were analysed by northern using probes detecting snR37, snR42 or snR190. Quantification of the northern data is presented in the histogram on the right. Ratios of immunoprecipitated snR190 versus immunoprecipitated snR37 (IP snR190/IP snR37) were obtained from phosphorimager scans of northern membranes. The resulting ratios were normalised to the ratio obtained with wild-type snR190, arbitrarily set at 1. Error bars correspond to standard deviations, computed from three independent biological replicates. Statistically significant differences determined using one-tailed Student’s t-test are indicated by asterisks (**= p<0.01; *= p<0.1; ns: not significant). (**B**) Extracts from the *NOP8::FPZ/snr190-[mut.C]* strains transformed with plasmids directing expression of wild-type snR190 (WT), the indicated snR190 mutants or the empty vector (ΔsnR190) were subjected to two consecutive ultracentrifugation steps. Nop8-FPZ was precipitated from these extracts with IgG Sepharose. Total proteins or RNAs were extracted from input extracts (INPUT (ultra)) or from immunoprecipitated samples (Nop8-FPZ IP) and analysed by western (top panel) or northern (bottom panel), respectively. Nop8-FPZ was detected using PAP, Npa1 and Dbp6 with specific antibodies. snR37 and snR190 were detected by northern using antisense oligonucleotide probes. Quantification of the northern data is presented in the histogram on the right. Ratios of immunoprecipitated snR190 versus immunoprecipitated snR37 (IP snR190/IP snR37) were obtained from phosphorimager scans of northern membranes. The resulting ratios were normalised to the ratio obtained with wild-type snR190, arbitrarily set at 1.

As Nop8 can interact directly with the snR190 snoRNP in absence of Npa1 complex formation, we asked whether the internal stem-loop of snR190 is also important for the interaction with Nop8. We transformed a *NOP8::FPZ/snr190-[mut.C]* strain with the plasmids directing expression of wild-type snR190, snR190-[mut.AB], snR190-[large Δstem] or the empty parental vector. Nop8-FPZ was then precipitated from extracts subjected to two consecutive ultracentrifugation steps to pellet pre-ribosomal particles. Northern analysis indicated that the co-precipitation efficiency of snR190-[large Δstem] was reduced 3-fold relative to wild-type snR190 (Figure 4B). Mutations of the A and B boxes led to an increase in snR190 co-precipitation, which is likely due to reduced association with pre-ribosomal particles (see Figure 5). We conclude that the Nop8-snR190 interaction involves major contacts with the internal stem-loop but other features of the snoRNP might also contribute to the interaction.

**Figure 5.**
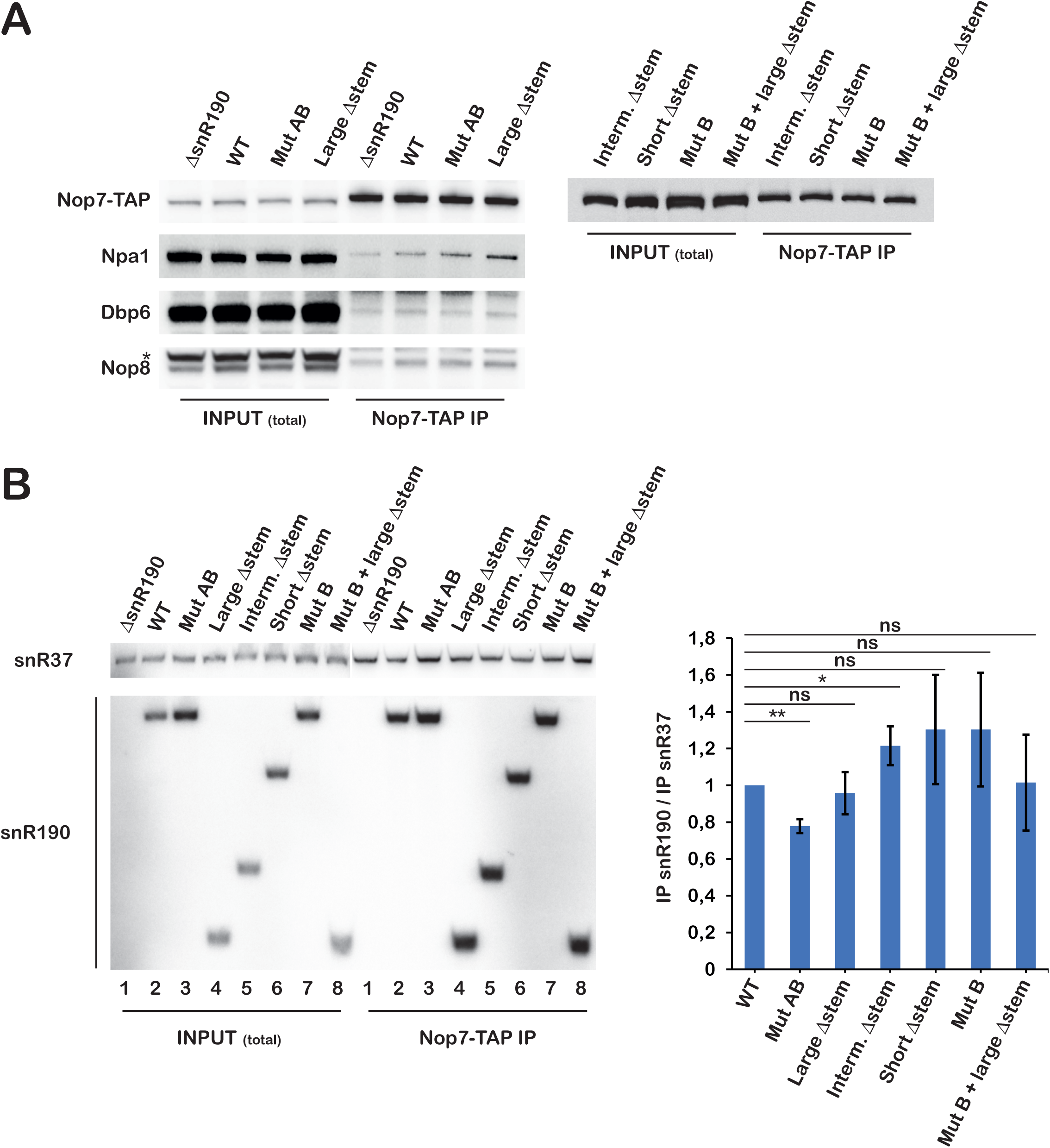
The internal stem-loop is not essential for snR190 incorporation or retention within pre-60S particles. Immunoprecipitation experiments were carried out with IgG Sepharose and extracts from *NOP7::TAP/snr190-[mut.C]* strains transformed with plasmids directing expression of wild-type snR190 (WT), the indicated snR190 mutants or the empty parental vector (ΔsnR190). Total proteins or RNAs were extracted from input extracts (INPUT (total)) or from immunoprecipitated samples (Nop7-TAP IP) and analysed by western (**A**) or northern (**B**), respectively. (**A**) Nop7-TAP was detected using PAP. For a subset of samples, Npa1, Dbp6 and Nop8 were also detected with specific antibodies. The star indicates an unknown polypeptide detected by the anti-Nop8 antibodies. (**B**) snR37 and snR190 were detected by northern using antisense oligonucleotide probes. Quantification of the northern data is presented in the histogram on the right. Ratios of immunoprecipitated snR190 versus immunoprecipitated snR37 (IP snR190/IP snR37) were obtained from phosphorimager scans of northern membranes. The resulting ratios were normalised to the ratio obtained with wild-type snR190, arbitrarily set at 1. Error bars correspond to standard deviations, computed from three independent biological replicates. Statistically significant differences determined using one-tailed Student’s t-test are indicated by asterisks (**= p<0.01; *= p<0.1; ns: not significant).

### The internal stem-loop of snR190 is not essential for snR190 association with pre-ribosomal particles

We next investigated the contribution of snR190 sequence elements to the stable integration of snR190 within pre-ribosomal particles. We transformed the plasmids directing expression of wild-type snR190 and the above-described snR190 mutants into a *snr190-[mut.C]* strain expressing TAP-tagged Nop7, to allow precipitation of 90S and pre-60S particles. These particles were precipitated with Nop7-TAP with the same efficiency in all cases, as assessed by the western analysis of precipitated Nop7-TAP (Figure 5A) and the northern analysis of precipitated pre-rRNAs (Supplementary Figure S10). Mutations of both A and B boxes of snR190 led to a 20% decrease in snR190 co-precipitation efficiency (Figure 5B). In contrast, the large internal stem-loop deletion or the combination of the large stem-loop deletion and the box B mutations did not affect snR190 co-precipitation efficiency (Figure 5B). These results suggest that the ability of snR190 to base pair with the pre-rRNA via either box A or B contributes to its stable integration within 90S/pre-60S pre-ribosomal particles but that its internal stem-loop structure has no influence. We also evaluated by western the co-precipitation of Npa1, Dbp6 and Nop8 in cells expressing snR190-[mut.AB] or snR190-[large Δstem], as well as cells expressing wild-type snR190 or lacking snR190 as controls (Figure 5A). Lack of snR190 internal stem-loop structure did not reduce the co-precipitation efficiency of Npa1, Dbp6 and Nop8. Altogether, these results suggest that the internal stem-loop structure of snR190 is required neither for the integration nor for the retention of snR190 and the Npa1 complex within pre-ribosomal particles.

### A strong interaction between the Npa1 complex and snR190 is important for efficient pre-60S particle maturation

Removal of the internal stem-loop structure of snR190 does not affect the association of snR190 or Npa1 complex members with pre-ribosomal particles. Yet it strongly weakens the association between Npa1 complex members and snR190. This decreased association may impact pre-60S particle maturation. To test this hypothesis, we analysed by northern the effects of snR190 mutations on pre-rRNA processing (Figure 6). We observed, as previously reported (16), that the absence of snR190 or the simultaneous mutation of both A and B boxes complementary to the pre-rRNA led to an increase in the levels of 27SA2 pre-rRNA and a decrease in 27SB pre-rRNA levels (Figure 6, lanes 1, 2 and 3; see also Supplementary Figure S1 for a scheme of pre-rRNA processing in *S. cerevisiae*). As a result, the 27SB/27SA2 ratio was halved when compared to the wild-type situation (Figure 6). The deletion of snR190 internal stem-loop also led to a similar, albeit less pronounced, pre-rRNA processing phenotype (Figure 6, lanes 4 and 8), resulting in a 27SB/27SA2 ratio diminished by about 30% compared to the wild-type control. In contrast, the box B mutations or the short and intermediate deletions within the internal stem-loop had no significant impact on pre-rRNA processing (Figure 6, lanes 5, 6 and 7). We also analysed the effects of these snR190 mutations in cells lacking both snR190 and snR37, the two major snoRNA components of early pre-60S particles (5), to determine whether the lack of snR37 would exacerbate the effects of snR190 mutations. In fact, the pre-rRNA processing defects elicited by the snR190 mutations were essentially identical in the *snr190-[mut.C]* and the *Δsnr37/snr190-[mut.C]* backgrounds (Supplementary Figure S11). We note that the deletion of snR190 internal stem-loop strongly weakens the direct interaction between snR190 and the Npa1 complex, while the box B mutations, or the short and intermediate internal stem deletions had no, or only a moderate effect on this interaction (Figure 4). Our data therefore suggest that a direct, high-affinity interaction between snR190 and the Npa1 complex within early pre-60S particles is required for their efficient maturation.

**Figure 6.**
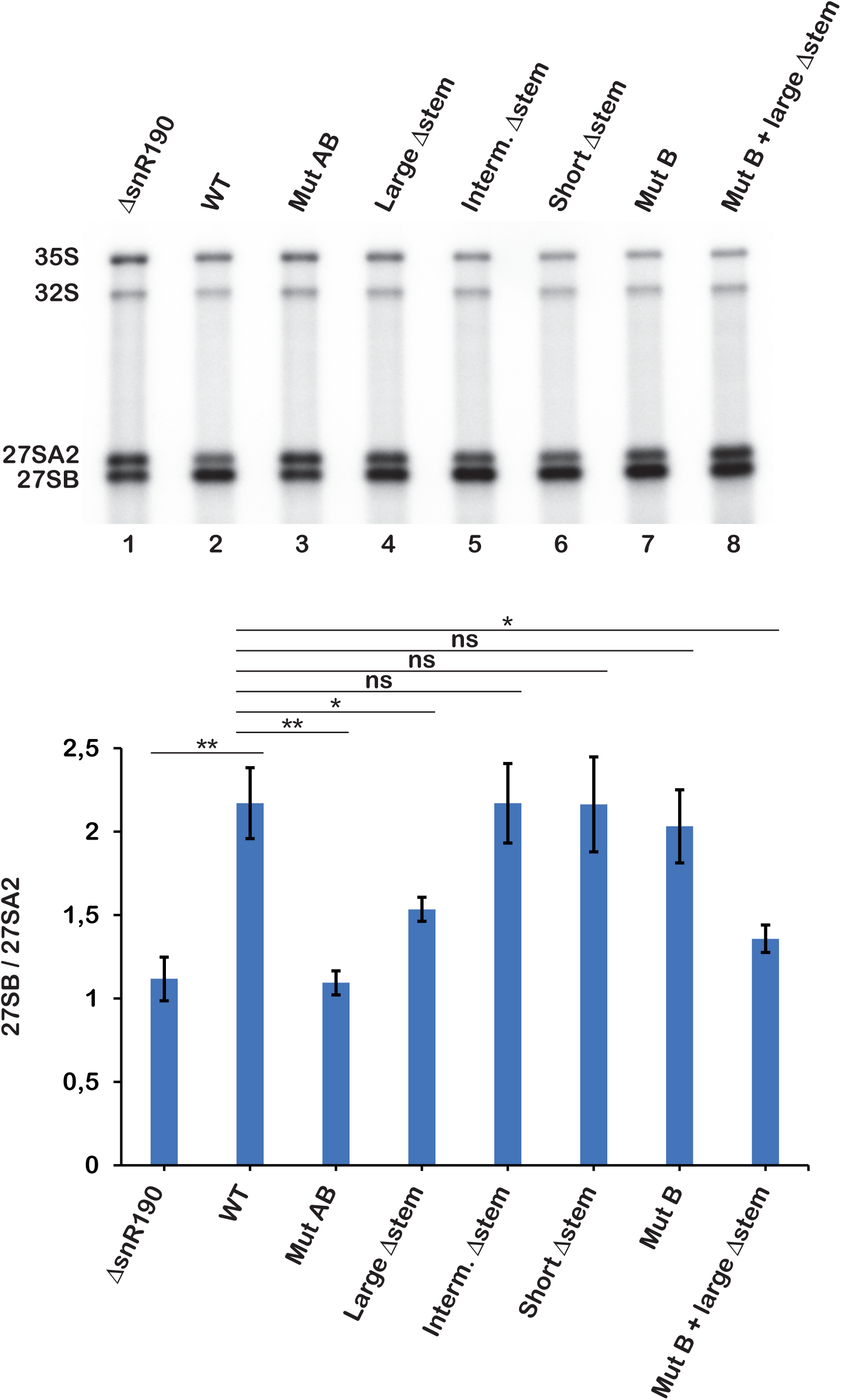
snR190 internal stem-loop is important for normal pre-rRNA processing. Total RNAs were extracted from *snr190-[mut.C]* strains transformed with plasmids directing expression of wild-type snR190 (WT), the indicated snR190 mutants or the empty parental vector (ΔsnR190) and analysed by northern. The indicated pre-rRNAs were detected with the rRNA2.1 probe. Quantification of the northern data is presented in the histogram below the northern. Levels of 27SA2 and 27SB pre-rRNAs were obtained from phosphorimager scans of northern membranes. Shown are the ratios of 27SB/27SA2 pre-rRNA levels. Error bars correspond to standard deviations computed from three independent biological replicates. Statistically significant differences determined using one-tailed Student’s t-test are indicated by asterisks (**= p<0.01; *= p<0.1; ns: not significant).

## Discussion

We have investigated the nature of the interactions between the Npa1 complex and the snR190 snoRNA, two components of the first pre-60S particles that may help promote the folding and/or clustering of the 5’ and 3’ domains of 25S rRNA. We previously showed that lack of snR190 weakens the association of Npa1 complex members with pre-ribosomal particles (16). Here we show that conversely, depletion of Npa1 or Nop8 severely weakens the association of snR190 with pre-ribosomal particles. The effect seems more drastic and specific to snR190 in the case of Nop8 depletion. This finding may reflect a direct interaction between Nop8 and the snR190 snoRNP that may help recruit this snoRNP into and/or tether it within pre-ribosomal particles. In contrast, Dbp6 depletion has little effect on snR190 association with pre-ribosomal particles, although we cannot rule out the establishment of improper snR190/pre-rRNA interactions when the catalytic activity of the Npa1 complex is missing. CRAC data previously indicated that Npa1 directly binds snR190 and that this snoRNA constitutes a major partner of Npa1 (10). We now demonstrate that all members of the Npa1 complex interact with snR190 outside pre-ribosomal particles, and therefore that a module constituted by the snR190 snoRNP bound to the Npa1 complex can be detected. Northern data and above all pCp labelling experiments strongly suggest that it is the only snoRNP that the Npa1 complex can interact with outside pre-ribosomal particles. To our knowledge, this is the first description of an independent macromolecular assembly involved in large-ribosomal-subunit synthesis constituted by a snoRNP chaperone bound to a defined protein complex. Another somewhat similar macromolecular assembly, containing the U3 snoRNP chaperone, nucleolin, Rrp5 and Dbp4, involved in small-ribosomal subunit synthesis in humans had been previously described (46). However, this assembly, whose full composition was not established, could only be detected when pre-rRNA transcription was blocked or when tUTP proteins were depleted.

Npa1 complex members are not required for normal accumulation of snR190 and hence are likely not snoRNP core proteins. snR190 is most probably stabilized by the canonical box C/D snoRNP proteins, consistent with the finding that box C mutations abolish snR190 accumulation (16). Moreover, it has been established that Nop58 and the methyltransferase Nop1, two core proteins of canonical box C/D snoRNPs, are components of the snR190 snoRNP (42,47). In accordance with these findings, we observed that Nop1 is co-purified with Rsa3 in the presence, but not in the absence of snR190. Conversely, lack of snR190 does not weaken the association between Rsa3 and its Npa1 complex members Npa1 and Dbp6. The absence of snR190 does not prevent the association between Nop8 and Rsa3 either, but strikingly, while Nop8 depletion has no effect on snR190 levels (see for example Figure 1B), lack of snR190 reduces Nop8 steady-state accumulation ((16), Figures 4B and 5A). This finding strengthens the hypothesis of a direct interaction between Nop8 and the snR190 snoRNP. One possible explanation is that binding of the snR190 snoRNP promotes/stabilizes Nop8 folding.

It remains unclear at present whether the snR190 snoRNP/Npa1 complex interaction is established prior to a common integration within pre-ribosomal particles or whether snR190 snoRNP and the Npa1 complex are released together from pre-ribosomal particles. Moreover, it is also possible that the snR190 snoRNP and the Npa1 complex may not associate prior to their integration into pre-ribosomal particles but are extracted together from these particles during the ultracentrifugation steps. However, the fact that the snR190 snoRNP/Npa1 complex module is also detected without ultracentrifugation of the extracts under conditions of RNA Pol I inhibition suggests that it exists in a free form in the cell. The interaction between the snR190 snoRNP and the Npa1 complex requires Npa1 and Nop8, but not Npa2. The requirement for Npa1 can be explained by the fact that this protein is essential for Npa1 complex formation and that it directly binds to snR190 according to CRAC data (10). Nop8 interacts directly with the free snR190 snoRNP in absence of Npa1 complex formation. It remains to be determined whether Nop8 solely binds to the snR190 snoRNA or whether it also interacts with snoRNP proteins. Interestingly in that respect, an interaction between Nop8 and Nop1, the methyltransferase component of box C/D snoRNPs, has been detected by a high-throughput cross-linking analysis (48). We initially hypothesized that Nop8 RRM, which is expected to bind RNA, could help tether the Npa1 complex to snR190 by a direct interaction with this snoRNA. However, our data demonstrate that the RRM is dispensable for Nop8 interaction with the free snR190 snoRNP. It contributes instead to the anchoring of Nop8 within pre-ribosomal particles, most probably via direct interaction(s) with the pre-rRNA.

The free snR190 snoRNP interacts with the Npa1 complex via, at least, Npa1 and Nop8. Npa1 establishes direct contacts with snR190 box B and with an internal stem-loop structure specific to snR190. Consistent with these binding sites, we found that removal of the entire stem-loop structure, and hence of two of the three Npa1 cross-linking regions, severely weakens the interaction outside pre-ribosomal particles between snR190 and the Npa1 complex purified via Rsa3. When we used Nop8 as bait, we also detected a reduced interaction with snR190, which was, however, far less diminished than when the purification was performed with Rsa3. We suppose that Nop8 can still interact to a significant extent with the snR190 snoRNP lacking the internal stem-loop structure, maybe via Nop1 (48). Nevertheless, our data clearly show that snR190 internal stem-loop is a very important contributor to the interaction between the Npa1 complex and the snR190 snoRNP, at least outside pre-ribosomal particles. This stem-loop structure has been conserved in snR190 homologues throughout Ascomycota, presumably due to its contribution to Npa1 complex binding (42). The fact that removal of this structure does not affect the accumulation of snR190 is consistent with the notion that Npa1 binding is not necessary for snR190 stability. We were somewhat surprised, however, that removal of the internal stem-loop structure of snR190 has no effect on snR190 association with pre-ribosomal particles, as assessed by immunoprecipitation experiments with tagged Nop7 as bait. This undiminished co-precipitation with Nop7 of snR190 lacking the internal stem-loop indicates that this structure is required neither for the integration, nor for the retention of snR190 within pre-ribosomal particles. The association of truncated snR190 with the pre-ribosomal particles may rely in part on the residual interaction with Nop8 and on base-pairing interactions with the pre-rRNA via box A. Indeed, the 20% reduction in co-precipitation efficiency with tagged Nop7 of snR190 featuring mutations within boxes A and B suggests that snoRNA/pre-rRNA base-pairing interactions contribute to the stability of snR190 association with pre-ribosomal particles. Our data also lead us to conclude that the removal of the internal stem-loop of snR190 does not affect the integration or retention within pre-ribosomal particles of Npa1, Nop8 and Dbp6.

Hence, both snR190 lacking the internal stem-loop structure and the Npa1 complex are present within pre-60S particles. Nevertheless, removal of this internal structure leads to a slightly aberrant pre-rRNA processing phenotype, characterized by reduced 27SB pre-rRNA levels relative to 27SA2 pre-rRNA levels. It is reminiscent of the defect induced by lack of snR190 (16), although it is milder. This processing phenotype may indicate impaired maturation of 27SA2-containing pre-60S particles and/or reduced stability of 27SB-containing pre-60S particles. Since the internal stem-loop structure of snR190 is required for high-affinity binding between the Npa1 complex and snR190 outside pre-ribosomal particles, it is tempting to propose that its deletion will also reduce the strength of the interactions between the Npa1 complex and snR190 within pre-60S particles and cause the pre-rRNA processing defect described above. A direct and strong interaction between the Npa1 complex and the snR190 snoRNP, mediated by its internal stem-loop structure, within the 27SA2-containing pre-60S particles may be required for snR190 and/or the Npa1 complex to correctly exert their role in pre-rRNA folding. This snR190 snoRNP/Npa1 complex interaction may correctly position the entire module relative to the pre-rRNA and/or the proteins present in their vicinity within the particles. For example, in 25S rRNA domain I, the Npa1-binding site and the sequence complementary to snR190 box B lie adjacent to the binding sites of Prp43, Rpl3 and Rrp5 (37,38). Available data suggest direct interactions between Prp43 and Npa1 and between Prp43 and Rrp5 (38,49) and indicate that Rpl3 is placed in the vicinity of Npa1 complex members Npa1, Nop8, Dbp6 and Rsa3 (14). In 25S rRNA domain V, Dbp6, Npa1 and Dbp7 binding sites and the sequence complementary to snR190 box A lie adjacent to each other in or close to the PTC (10,15,30). Importantly, the structural proximities are also corroborated by genetic interactions. Synthetic enhanced or synthetic lethal interactions have been obtained when combining inactivation/mutation of genes encoding Npa1 complex members and genes encoding Rpl3, Dbp7, Dbp9 and Rbp95 (11,14,50). Thus, incorrect interactions between the Npa1 complex and the snR190 snoRNP may have negative structural and functional impacts on the pre-60S particle neighbourhoods just described.

## Data availability

Data supporting the findings of this study are available from the corresponding authors upon reasonable request.

## Acknowledgements

We are grateful to B. Pertschy for the gift of the *Δsnr37* strain. We thank all members of the Henry/Henras team for helpful discussions. We are grateful to Mia-Latifah Maronat and Christine Maheu for expert technical assistance. We thank Augustin Le Mière for experimental help.

