## Supplementary material for "The snoRNP chaperone snR190 and the Npa1 complex form a macromomecular assembly required for 60S ribosomal subunit maturation"

#### Supplementary materials and methods

##### Plasmids

Centromeric plasmids based on pHA113 (1) directing expression of Nop8 or Nop8ΔRRM were obtained as follows. A plasmid directing expression of Nop8 tagged with two IgG-binding domains of *S. aureus* protein A was obtained by amplifying *NOP8* open reading frame using *S.*

*cerevisiae* genomic DNA with appropriate primers (Supplementary Table S1) followed by cloning in the BamHI site of pHA113 using the In-Fusion system (Clontech). The resulting plasmid, pHA113-NOP8-ZZ, was used as template to generate plasmids pHA113-NOP8 and pHA113-NOP8ΔRRM-ZZ using the In-Fusion system (Clontech) and appropriate primers (Supplementary Table S1). Finally, pHA113-NOP8ΔRRM-ZZ was used as template to generate plasmid pHA113-NOP8ΔRRM using the In-Fusion system (Clontech) and appropriate primers (Supplementary Table S1). The *NOP8* or *NOP8ΔRRM* open reading frames inserted in the resulting plasmids were entirely sequenced.

#### Yeast strains

A *GAL::HA-dbp6* strain was produced by transforming strain MW3628 (MATα, ura3-52, his3-Δ200, trp1-Δ63, leu2-Δ1) (2) with a PCR cassette obtained with plasmid pFA6a-kanMX6-PGAL1-3HA (3) and primers listed in Supplementary Table S2. Clones having integrated the kanMX6 resistance gene were selected on YP medium supplemented with 2% galactose and G418 (Gibco, 200 μg/ml final concentration). A *GAL::HA-dbp6/NOC1::FPZ* (Flag-PreScission cleavage site-tandem IgG-binding domains from *S. aureus* protein A) was produced by transforming the *GAL::HA-dbp6* strain described above with a PCR cassette obtained with plasmid pBS1479-NAT-2XFlag-PPX-ZZ and primers listed in Supplementary Table S2. Clones having integrated the nourseothricin resistance gene were selected on YP medium supplemented with 2% galactose and nourseothricin (Jena Bioscience, 80 μg/ml final concentration). A *Δsnr37/snr190-[mut.C]* strain was obtained by mating a *Δsnr37* (*SNR37* locus invalidated by nourseothricin resistance gene insertion, a gift of B. Pertschy) and a *snr190-[mut.C]* (mutated in box C, (4)) strain. The resulting diploids were sporulated and *Δsnr37/snr190-[mut.C]* haploids selected by screening for nourseothricin resistance and through sequencing of the *SNR190* locus.

#### Western analysis

Rsa3-Flag was detected with a HRP-conjugated anti-Flag mouse monoclonal antibody (FG4R, Invitrogen) used at 1000-fold dilution. Nop1 was detected with *Xenopus laevis* anti-fibrillarin serum (from rabbits) used at 270-fold dilution.

#### Supplementary data

**Supplementary Figure S1.** Pre-rRNA processing in *S. cerevisiae*. (**A**) rDNA organisation. (**B**) Pre-rRNA processing steps.

### Supplementary figure S1

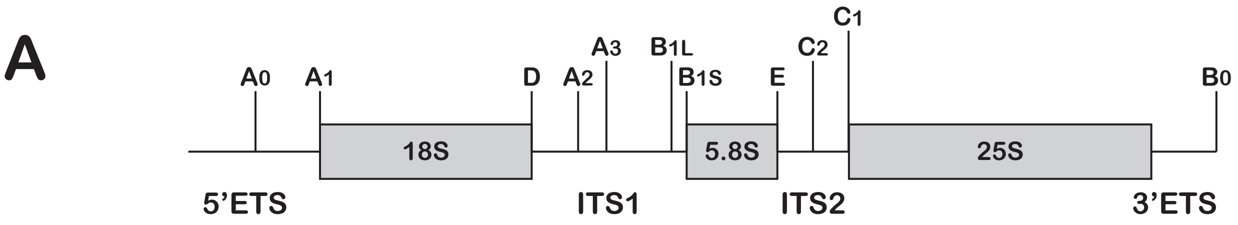

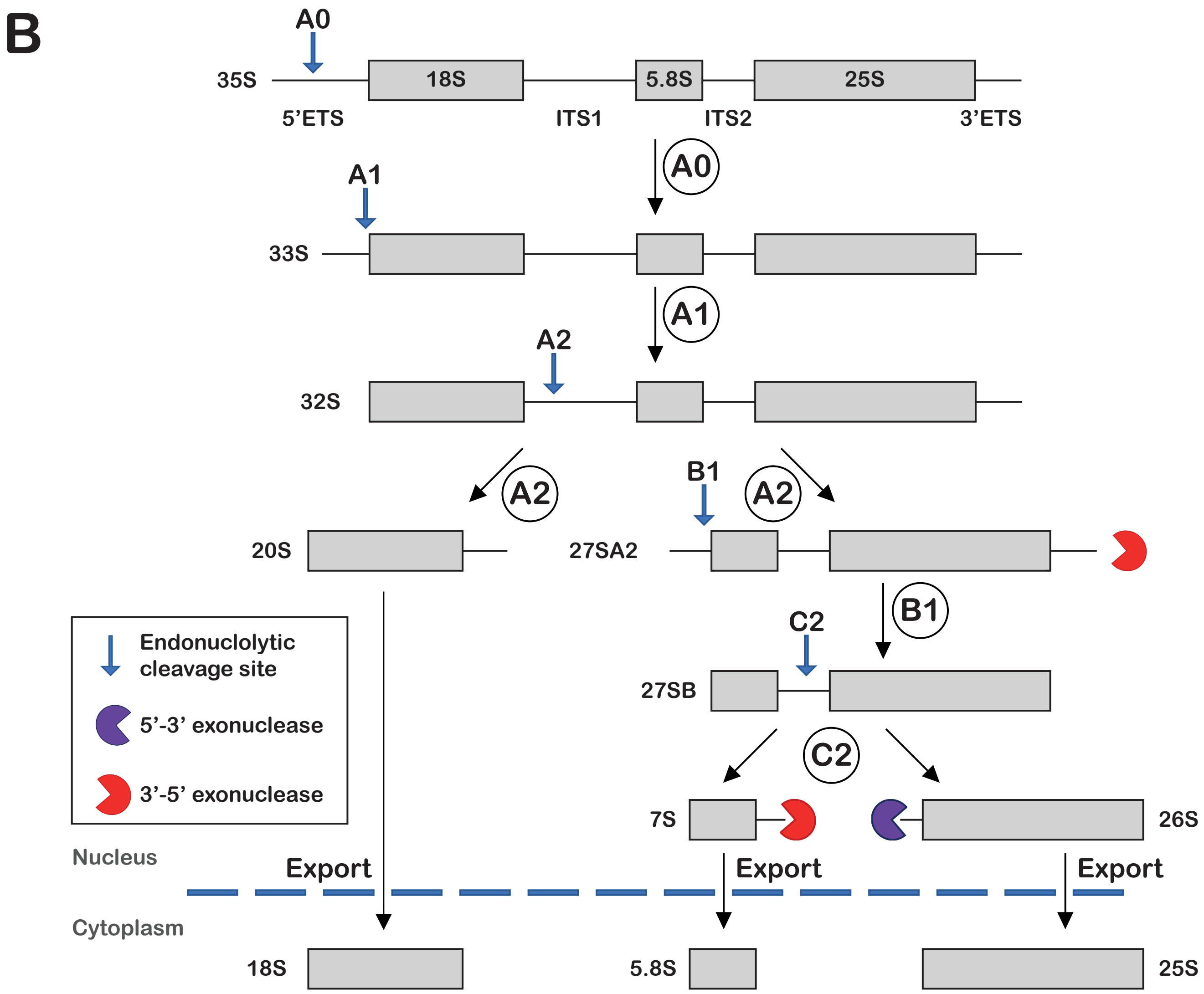

**Supplementary Figure S2.** Effects of Npa1 or Nop8 depletion on Noc1-FPZ association with pre-rRNAs. Immunoprecipitation experiments were carried out using IgG Sepharose and extracts from a BY4742 wild type strain (labelled Npa1 +, Noc1-FPZ - (**A**) or Nop8 +, Noc1-FPZ - (**B**)), a strain expressing Noc1-FPZ (labelled Npa1 +, Noc1-FPZ + (**A**) or Nop8 +, Noc1-FPZ + (**B**)), a *GAL::HA-npa1*/*NOC1::FPZ* strain (labelled Npa1 -, Noc1-FPZ + (**A**)) or a *GAL::HA-nop8*/*NOC1::FPZ* strain (labelled Nop8 -, Noc1-FPZ + (**B**)) expressing Noc1-FPZ and depleted of Npa1 or Nop8, respectively, following growth in glucose-containing medium for 14 hours. Total RNAs were extracted from input extracts (INPUT (total)) or from immunoprecipitated samples (IP) and analysed by northern. The indicated pre-rRNAs were detected using the 23S1 probe (**A**) or the rRNA2.1 probe (**B**).

### Supplementary figure S2

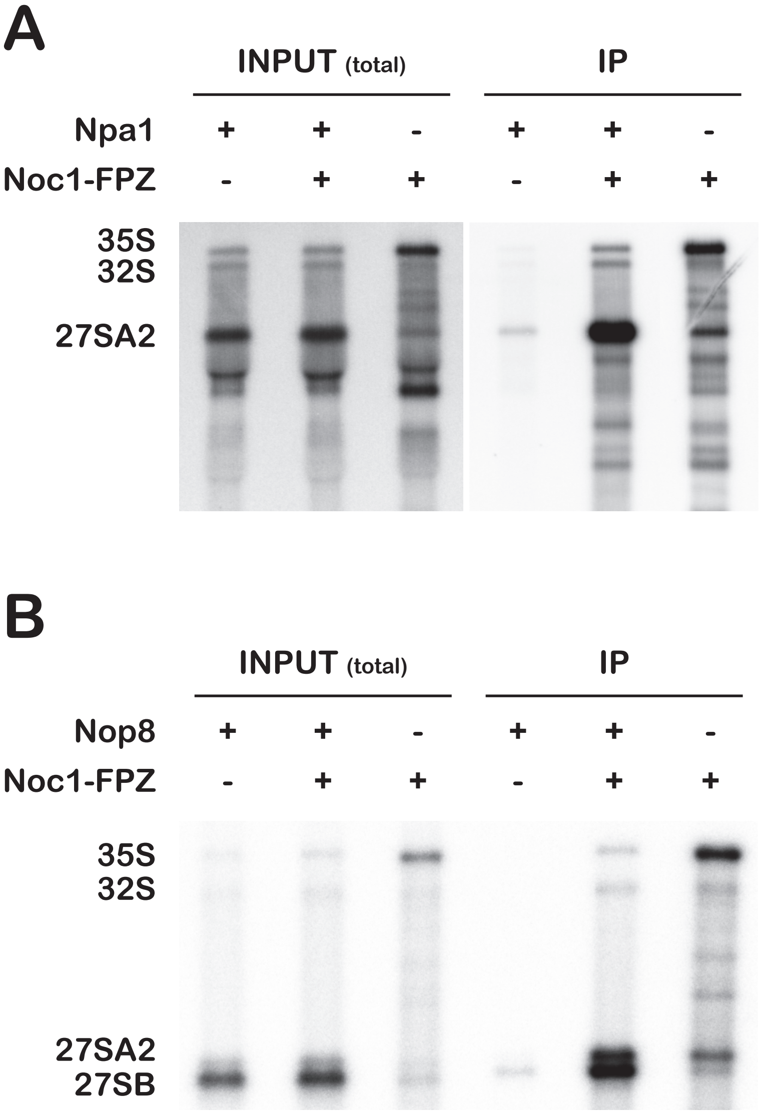

**Supplementary Figure S3.** Effects of Dbp6 depletion on Noc1-FPZ association with pre-rRNAs and on the efficiency of snoRNA co-precipitation with Noc1-FPZ. Immunoprecipitation experiments were carried out using IgG sepharose and extracts from a BY4742 wild-type strain (labelled Dbp6 +, Noc1-FPZ - (**C**)), a strain expressing Noc1-FPZ (labelled Dbp6 +, Noc1-FPZ +) and a *GAL::HA-dbp6*/*NOC1::FPZ* strain (labelled Dbp6 -, Noc1-FPZ +) expressing Noc1-FPZ and depleted of Dbp6 following growth in glucose-containing medium for 14 hours. Total proteins or RNAs were extracted from input extracts (INPUT (total)) or from immunoprecipitated samples (IP) and analysed by western (**A**) or northern for high (**B)** or low molecular weight RNA molecules **(C**). (**A**) Noc1-FPZ was detected using PAP, Dbp6 with specific antibodies. (**B**) The indicated pre-rRNAs were detected using the 23S1 probe. (**C**) The indicated snoRNAs were detected using antisense oligonucleotide probes. Quantification of northern data is presented in the histogram below the northern. Ratios of precipitated snoRNAs versus total snoRNAs present in the input extracts (IP/INPUT) were computed from phosphorimager scans of northern membranes. Ratios obtained for the *NOC1::FPZ* strain were arbitrarily set at 1.

### Supplementary figure S3

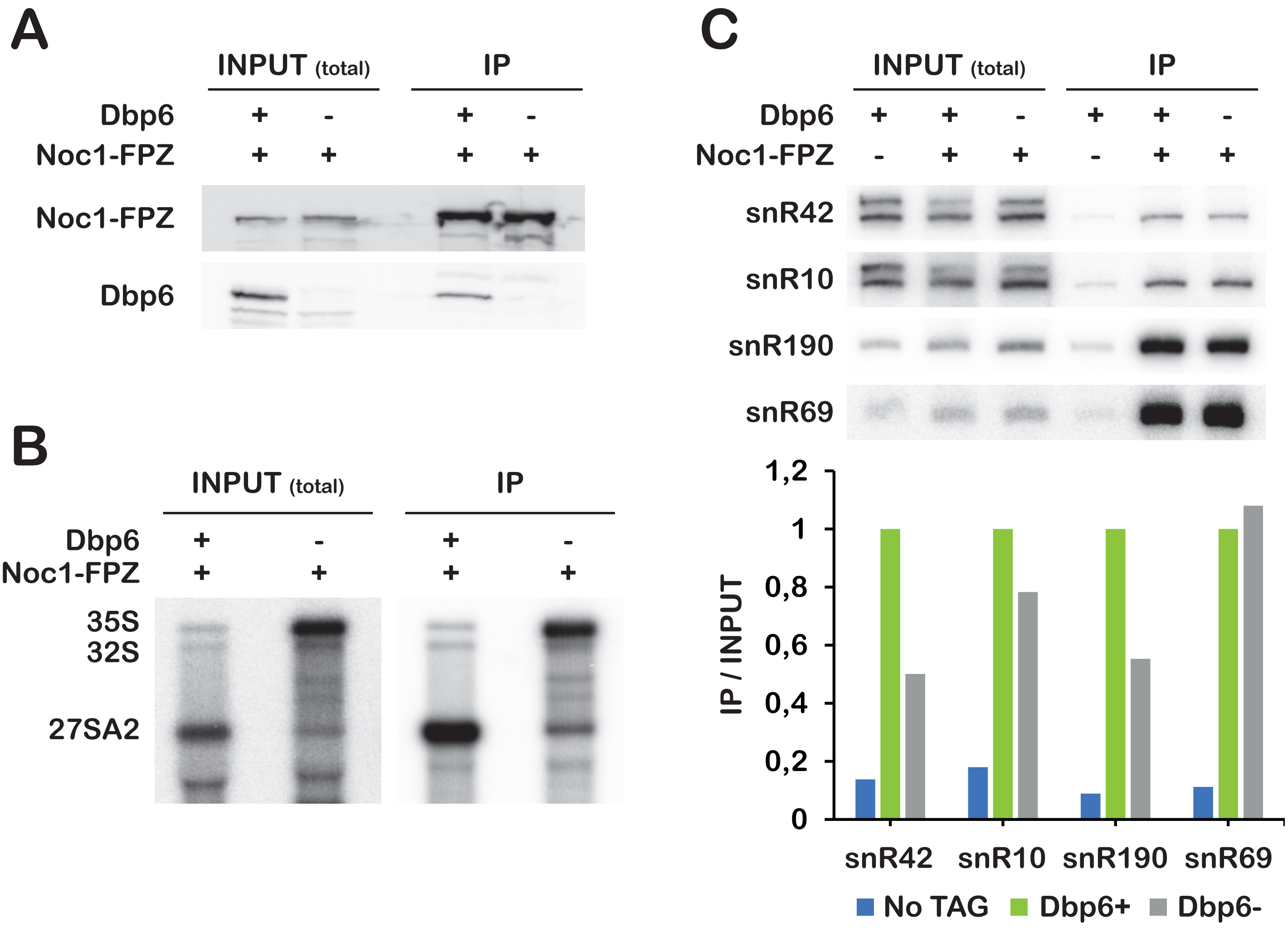

**Supplementary Figure S4.** Purification of the free Npa1 complex using a *rrn3.8/RSA3::FPZ* strain. (**A**) Northern analysis of pre-rRNA levels in the *rrn3.8/RSA3::FPZ* strain grown at 25°C (Pol I +) or transferred during 4 hours at 37°C (Pol I -) to inactivate RNA Pol I transcription. The indicated pre-rRNAs were detected using the 23S1 probe. (**B**) Western analysis of the Flag eluate of the tandem affinity purification performed with *rrn3.8/RSA3::FPZ* cells grown at 37°C for 4 hours. Rsa3-Flag was detected with anti-Flag-HRP antibodies, Npa1, Nop8 and Dbp6 with specific antibodies.

### Supplementary figure S4

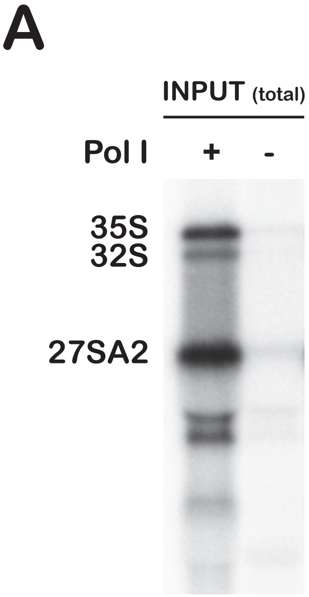

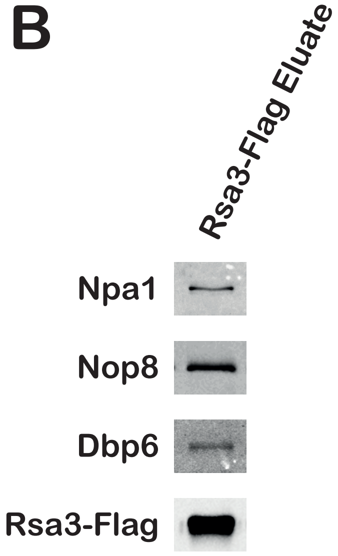

**Supplementary Figure S5.** Association of snR190 with free Npa1 complex in wild-type cells. Tandem affinity purification of Rsa3-FPZ was carried out with a wild-type strain extract subjected to two consecutive rounds of ultracentrifugation. (**A**) Northern analysis of pre-rRNAs extracted from the initial total clarified extract (INPUT (total)) and from the soluble fraction obtained after two consecutive ultracentrifugation steps (Ultra). The indicated pre-rRNAs have been detected using the 23S1 probe. (**B**) Western analysis of the eluate from the anti-Flag affinity column. Rsa3-Flag was detected with anti-Flag-HRP antibodies, Npa1, Dbp6 and Nop8 with specific antibodies. (**C**) Northern analysis of RNAs extracted from the soluble fraction obtained after two consecutive ultracentrifugation steps (Ultra) or the eluate from the anti-Flag affinity column (Rsa3-Flag Eluate), using oligonucleotide probes complementary to the indicated snoRNAs.

### Supplementary figure S5

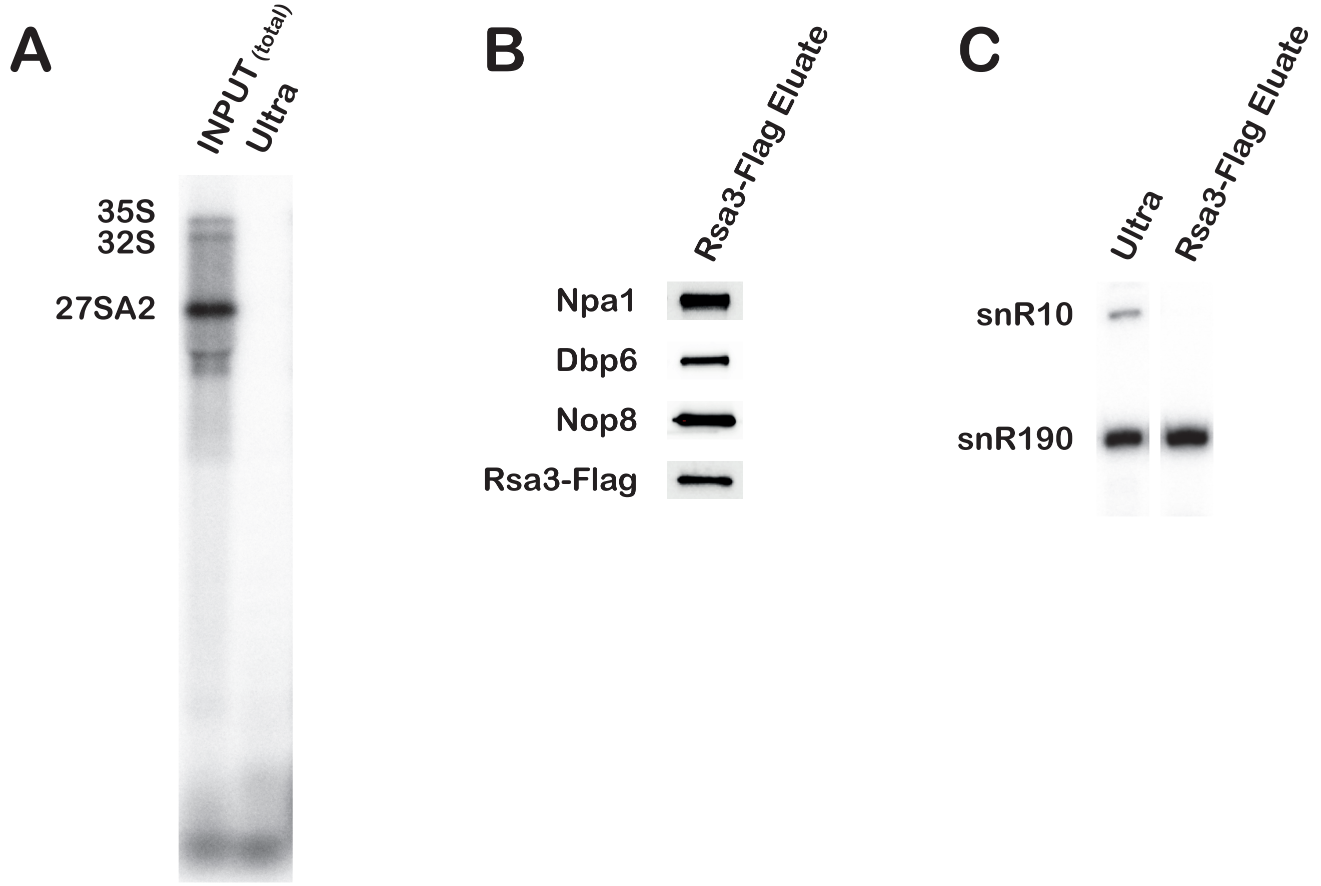

**Supplementary Figure S6.** Lack of snR190 does not prevent interactions between Rsa3 and its Npa1 complex partners. Tandem affinity purification of Rsa3-FPZ was carried out with *rrn3.8/RSA3::FPZ* (snR190 +) and *rrn3.8/snr190-[mut.C]/RSA3::FPZ* (snR190 -) strains grown in glucose-containing medium for 4 hours at 37°C to inactivate RNA Pol I transcription. Proteins in the initial clarified extracts (INPUT (*rrn3* ts)) and retained on the anti-Flag beads (Rsa3-Flag eluate) were analysed by western. (**A**) Rsa3-Flag was detected with anti-Flag-HRP antibodies, Npa1, Dbp6 and Nop8 with specific antibodies. The star indicates an unknown polypeptide detected by the anti-Nop8 antibodies. Quantification of western data is presented in the histogram below. Ratios of the quantity of proteins recovered in the Flag peptide eluate versus the quantity of Rsa3-Flag in the Flag peptide eluate (Flag eluate/Rsa3-Flag) were computed from Bioimager scans of western membranes. (**B**) Nop1 and Nhp2 were detected with specific antibodies.

### Supplementary figure S6

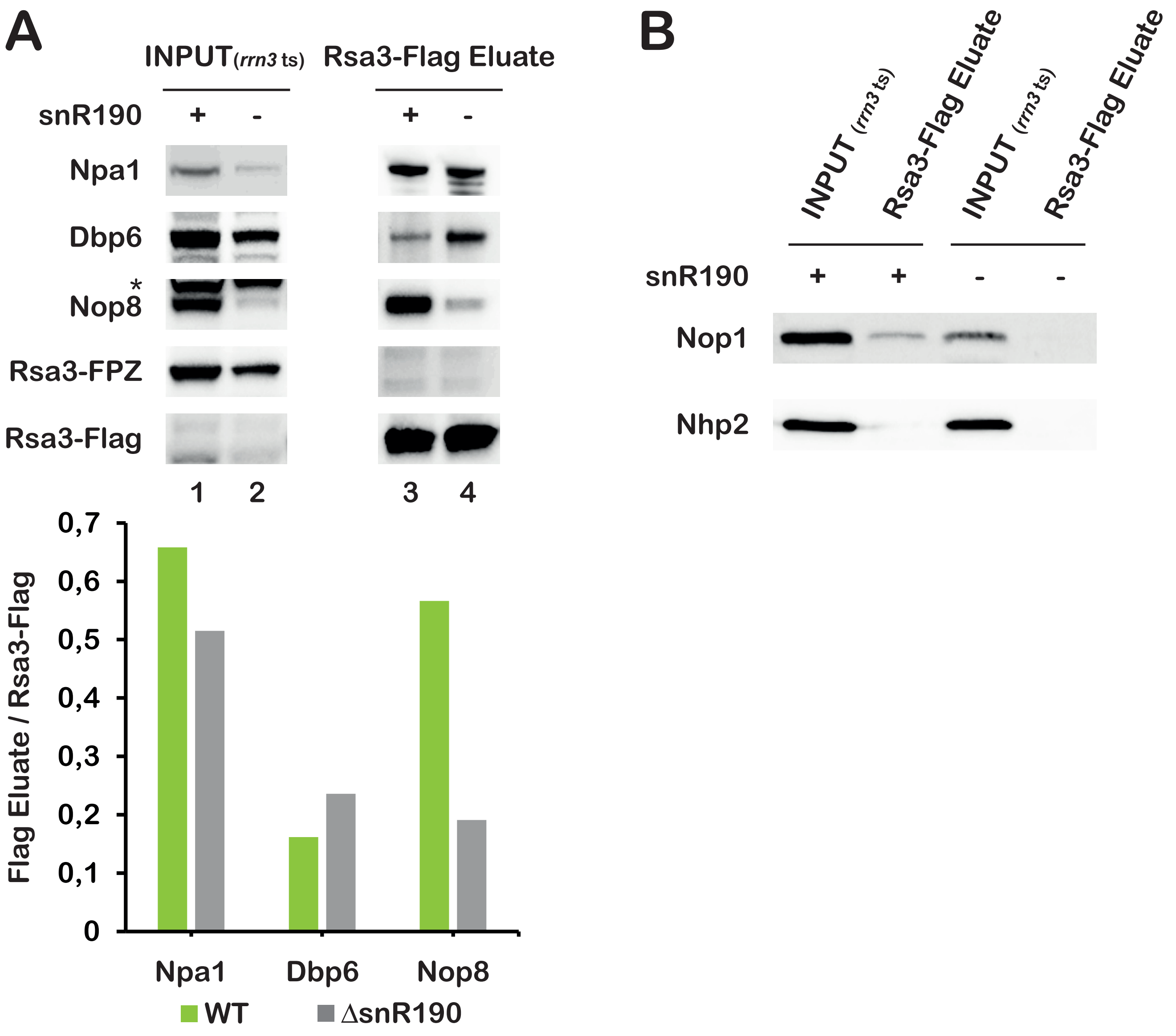

**Supplementary Figure S7.** (**A-C**) Tandem affinity purification of Rsa3-FPZ under conditions of Npa1, Npa2 or Nop8 depletion. (**A-B**) Tandem affinity purification of Rsa3-FPZ was carried out with *rrn3.8/GAL::HA-npa1/RSA3::FPZ* (**A**) and *rrn3.8/GAL::HA-npa2/RSA3::FPZ* (**B**) strains grown in glucose-containing medium for 8 hours at 37°C to inactivate RNA Pol I transcription and deplete Npa1 or Npa2, respectively. Proteins in the eluate fraction (Rsa3-Flag eluate) from the anti-Flag affinity column were analysed by western. Rsa3-Flag was detected with anti-Flag-HRP antibodies, Npa1, Dbp6 and Nop8 with specific antibodies. (**C**) Tandem affinity purification of Rsa3-FPZ was carried out with soluble extracts, obtained after two ultracentrifugation steps, from *RSA3::FPZ* (Nop8 +) or *GAL::HA-nop8/RSA3::FPZ* (Nop8 -) cells grown in glucose-containing medium for 14 hours. The indicated proteins in the eluate fraction (Rsa3-Flag Eluate) from the anti-Flag affinity column were analysed by western as described in (**A-B**). (**D**) Tandem affinity purification of Npa2-FPZ, Dbp6-FPZ or Nop8-FPZ under conditions of Npa1 depletion. Tandem affinity purification of Npa2-FPZ, Dbp6-FPZ or Nop8-FPZ was carried out with soluble extracts, obtained after two ultracentrifugation steps, from *NPA2::FPZ, DBP6::FPZ, NOP8::FPZ* (all labelled 'Npa1 +') or *GAL::HA-npa1/NPA2::FPZ, GAL::HA-npa1/DBP6::FPZ, GAL::HA-npa1/NOP8::FPZ* (all labelled 'Npa1 -') cells grown in glucose-containing medium for 14 hours. The indicated proteins in the eluate fraction (Flag Eluate) from the anti-Flag affinity columns were analysed by western as described in (**A-B**).

### Supplementary figure S7

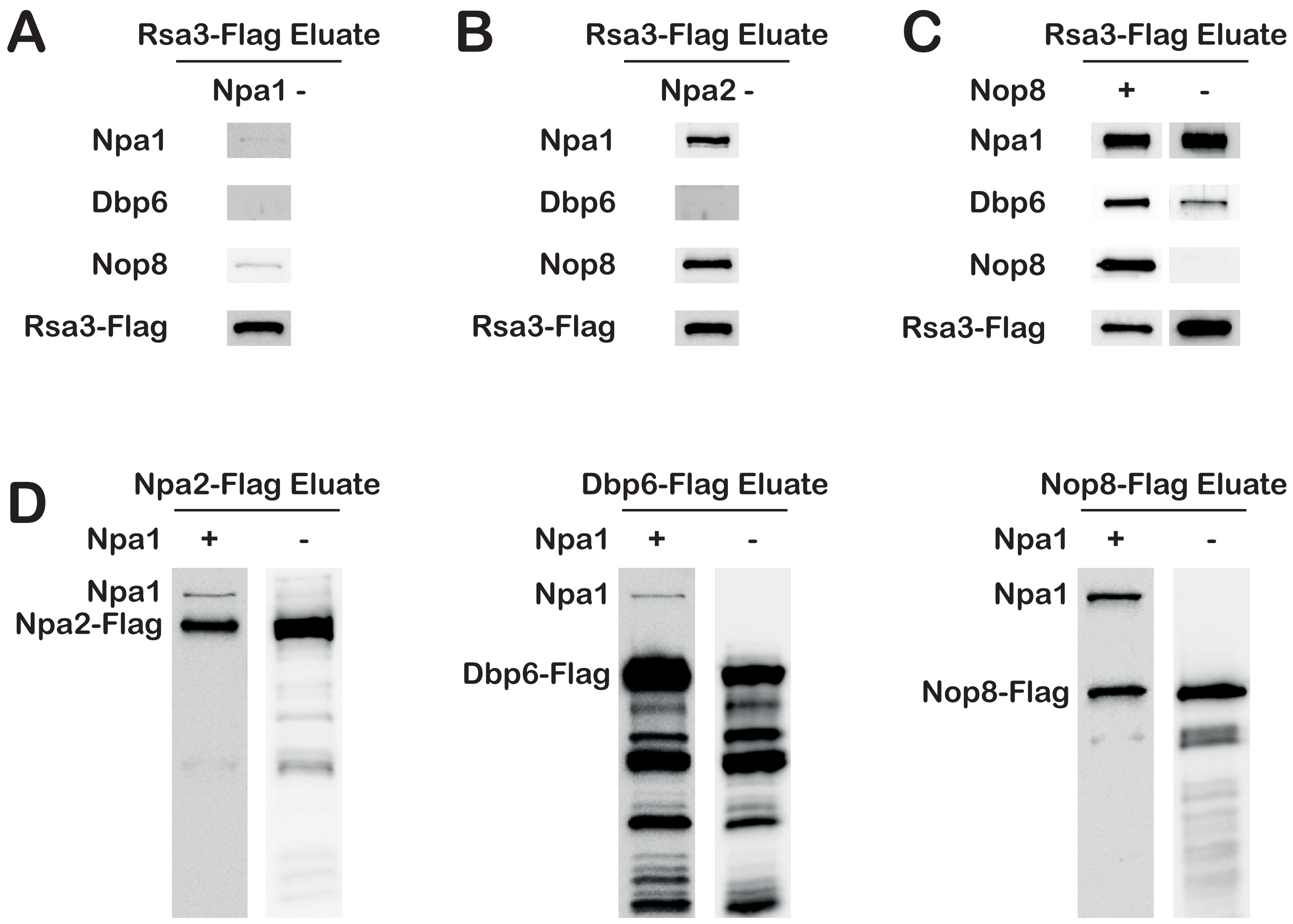

**Supplementary Figure S8.** Effects of Nop8 RRM deletion on snR190 association with pre-ribosomal particles and on pre-rRNA processing. (**A-B**) Immunoprecipitation experiments were carried out using IgG Sepharose and extracts from *GAL::HA-nop8/NOC1::FPZ* cells grown in in glucose-containing medium for 14 hours to deplete genome-encoded Nop8 and transformed with an empty parental vector (E.V.), or plasmids directing expression of wild-type Nop8 or Nop8ΔRRM. Total proteins or RNAs were extracted from input extracts (INPUT (total)) or from immunoprecipitated samples (Noc1-FPZ IP) and analysed by western (**A**) or northern (**B**). (**A**) Noc1-FPZ was detected using PAP, Npa1 and Nop8 with specific antibodies. The star highlights an unknown polypeptide detected by the anti-Nop8 serum. (**B**) snR37 and snR190 were detected with specific antisense oligonucleotide probes. Quantification of northern data is presented in the histogram on the right. Ratios of precipitated snoRNAs versus snoRNAs present in the input extracts (IP/INPUT) were computed from phosphorimager scans of northern membranes. Ratios obtained for the NOP8-expressing strain were arbitrarily set at 1. Error bars correspond to standard deviations computed from two technical replicates. (**C**) Total RNAs were extracted from the parental W303 wild-type strain, a strain expressing Nop8-HTP or Nop8ΔRRM-HTP and analysed by northern. The indicated pre-rRNAs were detected using the rRNA2.1 probe. Quantification of northern data is presented in the histogram on the right. Levels of 27SA2 and 27SB pre-rRNAs were obtained from phosphorimager scans of northern membranes. Shown are the ratios of 27SB/27SA2 pre-rRNA levels. Error bars correspond to standard deviations computed from three independent biological replicates.

### Supplementary figure S8

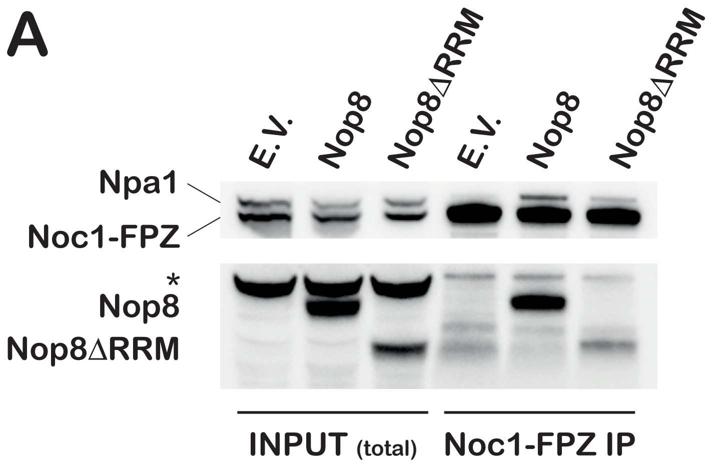

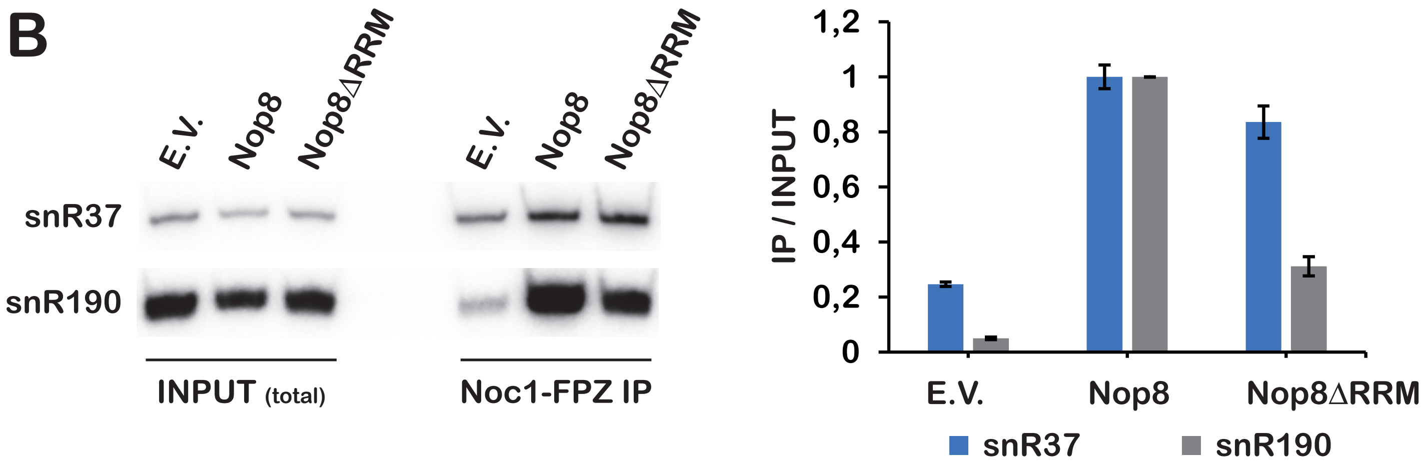

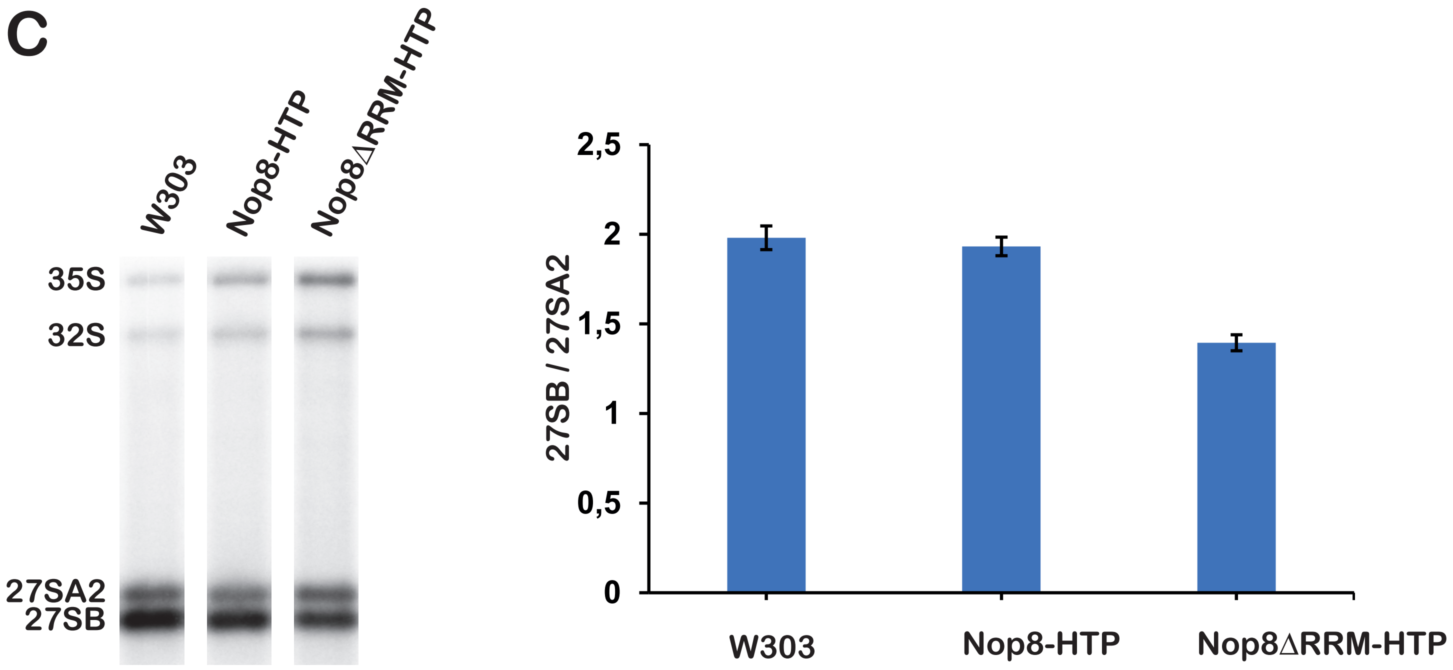

**Supplementary Figure S9.** (**A**) Secondary structure prediction of snR190 with the positions of Npa1 cross-linking sites identified by CRAC. The sequence of the mutated box B of the snR190-[mut.B] mutant is indicated. The segments of the internal stem-loop removed in the various truncation mutants are indicated by brackets. (**B**) Steady-state accumulation of snR190 mutants. Total RNAs were extracted from *snr190-[mut.C]* strains transformed with plasmids directing expression of wild-type snR190 (WT), the indicated snR190 mutants or the empty parental vector (ΔsnR190) and analysed by northern. snR37 and snR190 were detected with antisense oligonucleotide probes. Quantification of northern data is presented in the histogram on the right. Levels of snR37 and snR190 were obtained from phosphorimager scans of northern membranes. Shown are the ratios of snR190 over snR37 levels, normalised to the ratio obtained for wild-type snR190, arbitrarily set at 1. Error bars correspond to standard deviations computed from three independent biological replicates.

### Supplementary figure S9

**A**

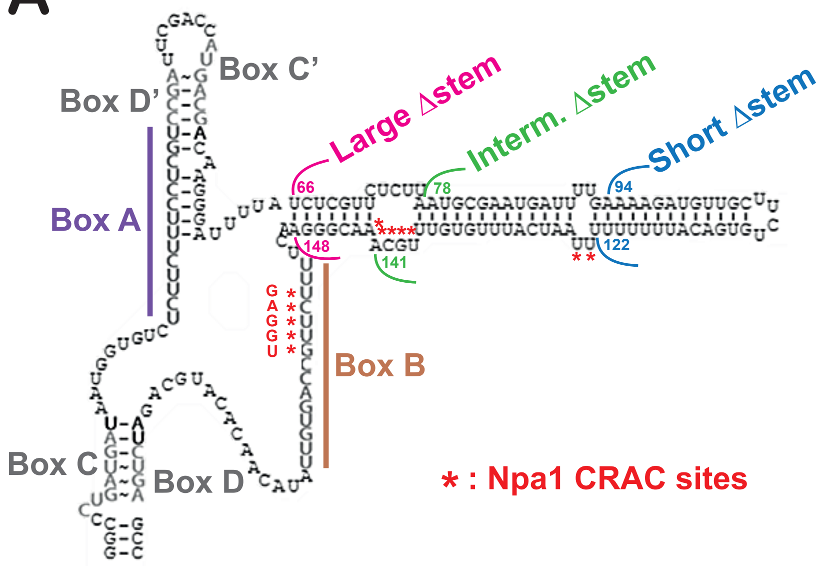

**B**

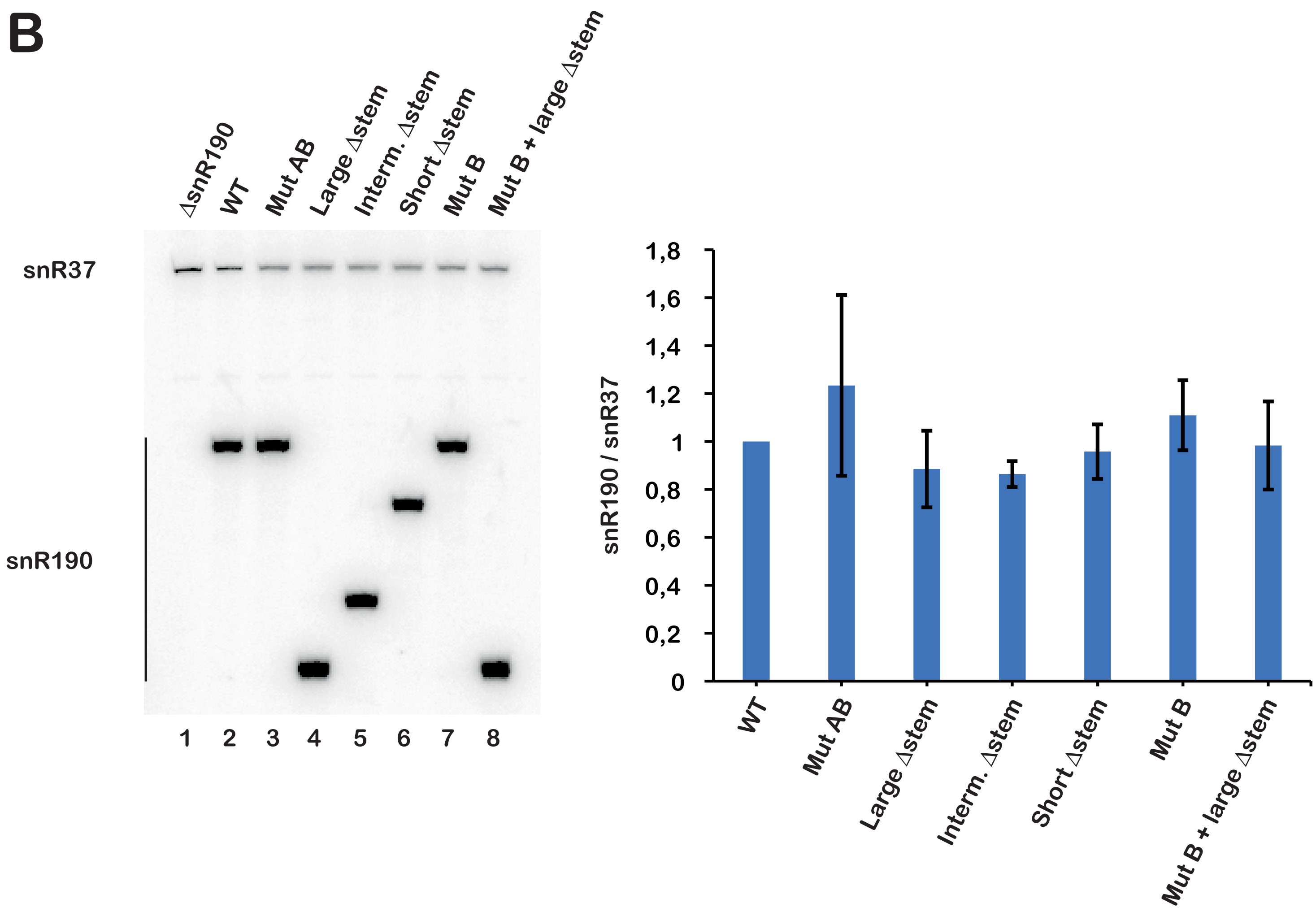

**Supplementary Figure S10.** Immunoprecipitation experiments were carried out with IgG Sepharose and extracts from *NOP7::TAP/snr190-[mut.C]* strains transformed with plasmids directing expression of wild-type snR190 (WT), the indicated snR190 mutants or the empty parental vector (ΔsnR190). The indicated pre-rRNAs extracted from the total extracts (INPUT (total)) or immunoprecipitated samples (Nop7-TAP IP) were detected by northern using the 23S1 probe.

### Supplementary figure S10

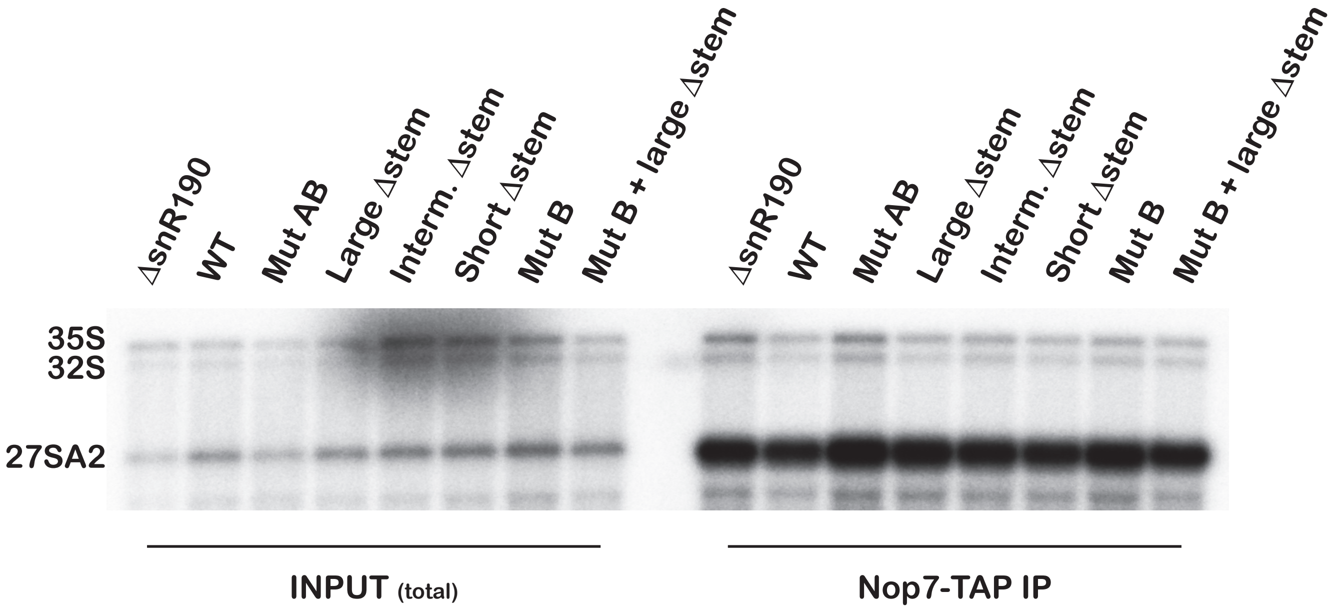

**Supplementary Figure S11.** Effects of snR190 mutants on pre-rRNA processing in *Δsnr37/snr190-[mut.C]* background. Total RNAs were extracted from *Δsnr37/snr190-[mut.C]* strains transformed with plasmids directing expression of wild-type snR190 (WT), the indicated snR190 mutants or the empty parental vector (ΔsnR190) and analysed by northern. (**A**) Analysis of pre-rRNA levels. The indicated pre-rRNAs were detected with the rRNA2.1 probe. Quantification of pre-rRNA northern data is shown in the histogram on the right. Levels of 27SA2 and 27SB pre-rRNAs were obtained from phosphorimager scans of northern membranes. Shown are the ratios of 27SB/27SA2 pre-rRNA levels. Error bars correspond to standard deviations computed from three independent biological replicates. Statistically significant differences determined using one-tailed Student's t-test are indicated by asterisks (***= $p<0.001$; **= $p<0.01$; *= $p<0.1$; ns: not significant). (**B**) Analysis of wild-type and mutant snR190 levels. snR10 and snR190 were detected with antisense oligonucleotide probes. Quantification of snoRNA northern data is shown in the histogram on the right. Levels of snR10 and snR190 were obtained from phosphorimager scans of northern membranes. Shown are the ratios of snR190 over snR10 levels, normalised to the ratio obtained for wild-type snR190, arbitrarily set at 1. Error bars correspond to standard deviations computed from three independent biological replicates.

### Supplementary figure S11

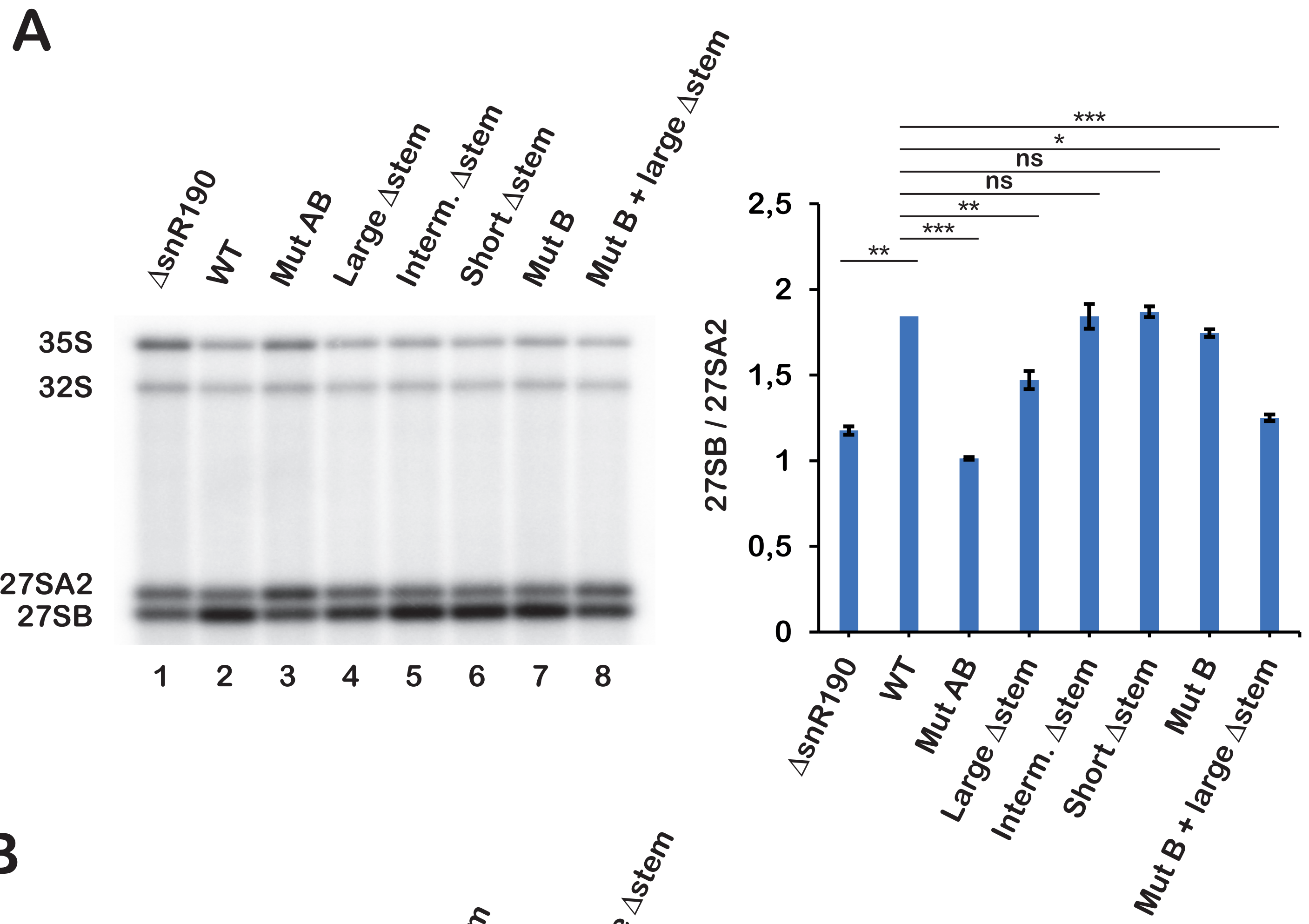

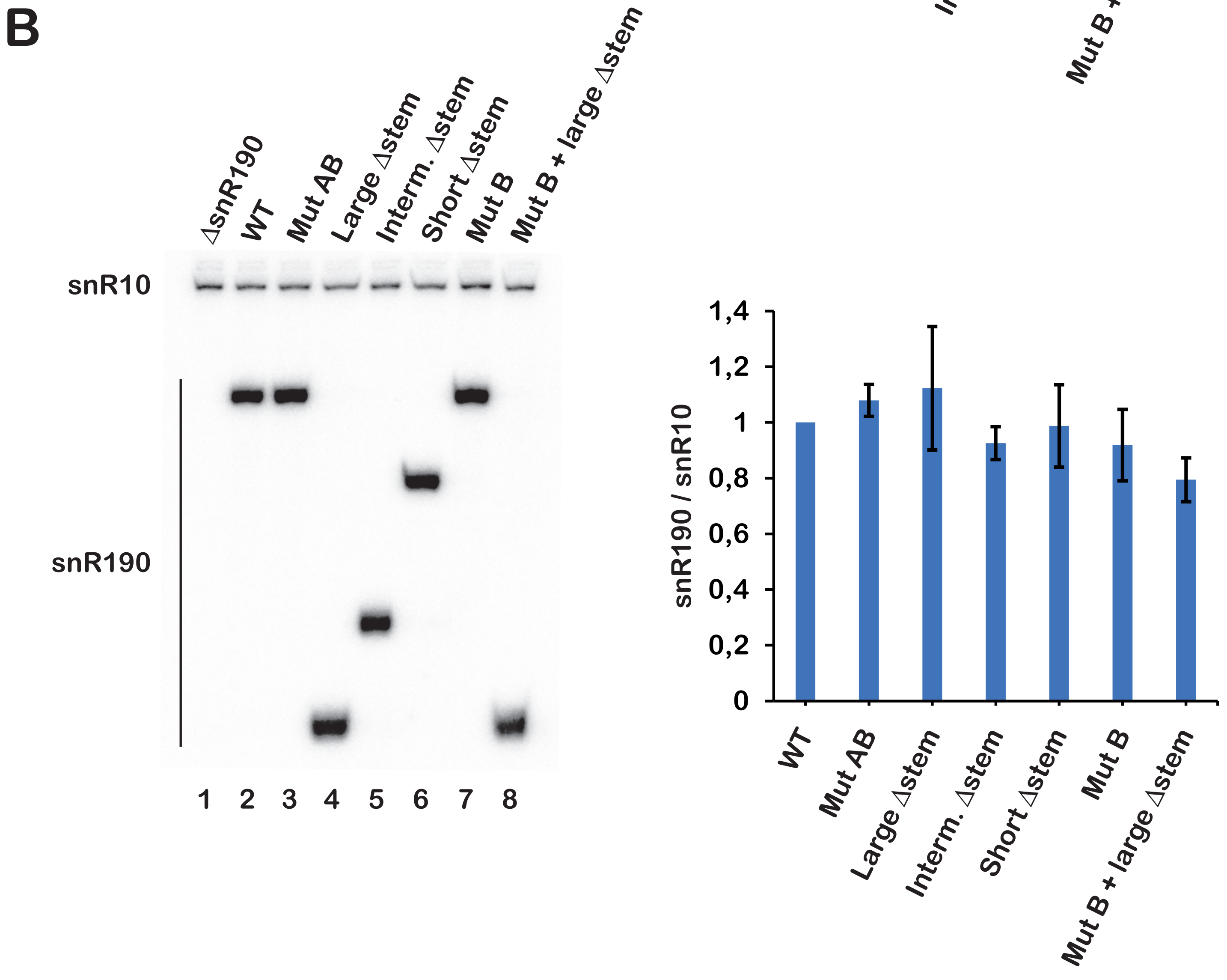

**Supplementary Table S1.** Primers used for plasmid construction.

| Plasmid | Primers used (5'-3') |
|---|---|
| snR190-[short Δstem] | ATGATTAACAAAATCATTCGCATTAAGAGAACG |
| | GATTTTGTTAATCATTTGTGTTTGCAAACGGG |
| snR190-[intermediate Δstem] | TTCCCGTTAAGAGAACGAGATAAAATCCCTTG |
| | TTCTCTTAACGGGAACTTTTCTTGCCAG |
| snR190-[large Δstem] | AGAAAAGTTAAAATCCCTTGTCGTCATGGTCG |
| | GATTTTAACTTTTCTTGCCAGTGTTATACAAC |
| snR190-[mut.B] | AACTTTGAGGTCCAGTGTTATACAACACATGCAGA |
| | CTGGACCTCAAAGTTCCCGTTTGCAAACACA |
| snR190-[mut.B-large Δstem] | AACTTTGAGGTCCAGTGTTATACAACACATGCAGA |
| | CTGGACCTCAAAGTTAAAATCCCTTGTCGTCATG |
| pHA113-NOP8-ZZ | GTTTCAGAGGGGATCCGATGGATAGTGTAATTCAAAAAAG |
| | GCATGCTGAAGGATCTAGAAGAAGCCCGCTCTTTG |
| pHA113-NOP8 | TTCTATAGTAGGATCCTTCAGCATGCCTTG |
| | GATCCTACTATAGAAGAAGCCCGCTCTTTG |
| pHA113-NOP8ΔRRM-ZZ | GGGATCCGAAGCCGAATTGGGAAAGCAC |
| | TCGGCTTCGGATCCCCTCTGAAACTC |
| pHA113-NOP8ΔRRM | TTCTATAGTAGGATCCTTCAGCATGCCTTG |
| | GATCCTACTATAGAAGAAGCCCGCTCTTTG |

**Supplementary Table S2.** Primers used for yeast strain construction.

| **Strain** | **Primers used (5'-3')** |
|---|---|
| NOC1::FPZ | Forward primer<br>TTGCATCTGCCGACGATTATGCTCAATATTTAGATCAAGAT<br>TCAGACTCCATGGACTACAAGGACGACGATG |
| | Reverse primer<br>ATTGTACAAAATTCTGTTTTCCTTATTTAATTTACAACA<br>CCGAAGTGTTTTACGACTCACTATAGGGCGAATTG |
| NOP8::FPZ | Forward primer<br>GATGCATTGAAGCACCGTAAGAGGAAACAATCAAAGAGCGGGCT<br>TCTTCTAATGGACTACAAGGACGACG |
| | Reverse primer<br>TGCAGCAGTGGAAATGGCTGTTAAGTTCGGTAAAAATGCTTTTTG<br>AATAAGCGCAGAGAAATCTATACTATATATGGCGTAATACGACTC<br>ACTATAGGG |
| NOP7::TAP | Forward primer<br>CACTCATCAAATTGTTGACAG |
| | Reverse primer<br>GAAGTGTTTCAATGCAAACG |
| GAL::HA-npa1 | Forward primer<br>TGATTGGATATTATTTCTCTAATCTATGCGGTACTACTT<br>CATCTAACAGAGAATTCGAGCTCGTTTAAAC |
| | Reverse primer<br>TTCTCCCTTCTCTGGTCGCGAGATCCATAGGCTTCGCTA<br>TGATTACTCATGCACTGAGCAGCGTAATCTG |
| GAL::HA-npa2 | Forward primer<br>TCTGTCTAAGATTTAGCTTGCCATCAATTATCTTTGGAA<br>AAACAGAGAGTGAATTCGAGCTCGTTTAAAC |
| | Reverse primer<br>AAATCTTGGGCATTGTCTGGGATAGATAGTTCTTCTGTAA<br>GATCACCCATGCACTGAGCAGCGTAATCTG |

| | |
|---|---|
| GAL::HA-nop8 | Forward primer TAACAAAGTATACAATAGGCCCATATCATTTTAGATTGTACCTGAAGTGAGAATTCGAGCTCGTTTAAAC |
| | Reverse primer TTATGGAAAATATTTCCGACAAAAATTCTTTTTGAATTACACTATCCATGCACTGAGCAGCGTAATCTG |
| GAL::HA-dbp6 | Forward primer TAGAAAGCATAACTGGCGATGAGATGTGCTCGAAACTGGACACCTTTACCGAATTCGAGCTCGTTTAAAC |
| | Reverse primer CTAGCTGCAGGAGCAGTCAATTGGCTAGGGTCAAATCTCGATGCAAACATGCACTGAGCAGCGTAATCTG |

**Supplementary Table S3.** Antisense oligonucleotides used as northern probes.

| | **Sequence (5'-3')** |
|---|---|
| Anti-snR3 | CGAATAAGACCGAGTGTTCA |
| Anti-snR39b | TATCAGCCCCGAAAGATGGTGTAATTT |

**Supplementary Table S4.** Inventory of the yeast strains used in this study with their genetic background, the enclosed plasmids and genotypes.

| **STRAIN** | **BACKGROUND** | **PLASMID** | **GENOTYPE** |
|---|---|---|---|
| BY4742 | - | - | *MATα, his3Δ1, leu2Δ0, lys2Δ0, ura3Δ0* |
| *RSA3::FPZ* | BY4742 | - | *MATα, his3Δ1, leu2Δ0, lys2Δ0, ura3Δ0, RSA3::FPZ (URA3)* |
| *DBP6::FPZ* | BY4742 | - | *MATα, his3Δ1, leu2Δ0, lys2Δ0, ura3Δ0, DBP6::FPZ (URA3)* |
| *NPA2::FPZ* | BY4742 | - | *MATα, his3Δ1, leu2Δ0, lys2Δ0, ura3Δ0, NPA2::FPZ (URA3)* |
| *NOP8::FPZ* | BY4742 | - | *MATα, his3Δ1, leu2Δ0, lys2Δ0, ura3Δ0, NOP8::FPZ (URA3)* |
| *GAL::HA-npa1/ RSA3::FPZ* | BY4742 | - | *MATα, his3Δ1, leu2Δ0, lys2Δ0, ura3Δ0, GAL::HA-npa1 (kanMX6), RSA3::FPZ (URA3)* |
| *GAL::HA-npa1/ DBP6::FPZ* | BY4742 | - | *MATα, his3Δ1, leu2Δ0, lys2Δ0, ura3Δ0, GAL::HA-npa1 (kanMX6), DBP6::FPZ (URA3)* |
| *GAL::HA-npa1/ NPA2::FPZ* | BY4742 | - | *MATα, his3Δ1, leu2Δ0, lys2Δ0, ura3Δ0, GAL::HA-npa1 (kanMX6), NPA2::FPZ (URA3)* |
| *GAL::HA-npa1/ NOP8::FPZ* | BY4742 | - | *MATα, his3Δ1, leu2Δ0, lys2Δ0, ura3Δ0, GAL::HA-npa1 (kanMX6), NOP8::FPZ (URA3)* |
| *GAL::HA-nop8/ RSA3::FPZ* | BY4742 | - | *MATα, his3Δ1, leu2Δ0, lys2Δ0, ura3Δ0, GAL::HA-nop8 (kanMX6), RSA3::FPZ (URA3)* |
| *NOC1::FPZ* | BY4742 | - | *MATα, his3Δ1, leu2Δ0, lys2Δ0, ura3Δ0, NOC1::FPZ (NAT)* |
| *GAL::HA-npa1/ NOC1::FPZ* | BY4742 | - | *MATα, his3Δ1, leu2Δ0, lys2Δ0, ura3Δ0, GAL::HA-npa1 (kanMX6), NOC1::FPZ (NAT)* |
| *GAL::HA-nop8/ NOC1::FPZ* | BY4742 | - | *MATα, his3Δ1, leu2Δ0, lys2Δ0, ura3Δ0, GAL::HA-nop8 (kanMX6), NOC1::FPZ (NAT)* |

| | | | |
|---|---|---|---|
| *GAL::HA-nop8/ NOC1::FPZ* | BY4742 | pHA113 | *MATα*, *his3Δ1*, *leu2Δ0*, *lys2Δ0*, *ura3Δ0*, *GAL::HA-nop8 (kanMX6)*, *NOC1::FPZ (NAT)* |
| *GAL::HA-nop8/ NOC1::FPZ* | BY4742 | pHA113-NOP8-ZZ | *MATα*, *his3Δ1*, *leu2Δ0*, *lys2Δ0*, *ura3Δ0*, *GAL::HA-nop8 (kanMX6)*, *NOC1::FPZ (NAT)* |
| *GAL::HA-nop8/ NOC1::FPZ* | BY4742 | pHA113-NOP8ΔRRM-ZZ | *MATα*, *his3Δ1*, *leu2Δ0*, *lys2Δ0*, *ura3Δ0*, *GAL::HA-nop8 (kanMX6)*, *NOC1::FPZ (NAT)* |
| *rrn3.8* | *rrn3.8* | - | *ade5*, *his7-2*, *leu2-112*, *trp1-289*, *ura3-52*, *rrn3.8* |
| *rrn3.8/RSA3::FPZ* | *rrn3.8* | - | *ade5*, *his7-2*, *leu2-112*, *trp1-289*, *ura3-52*, *rrn3.8, RSA3::FPZ (NAT)* |
| *rrn3.8/GAL::HA-npa1/RSA3::FPZ* | *rrn3.8* | - | *ade5*, *his7-2*, *leu2-112*, *trp1-289*, *ura3-52*, *rrn3.8, GAL::HA-npa1 (kanMX6), RSA3::FPZ (NAT)* |
| *rrn3.8/GAL::HA-npa2/RSA3::FPZ* | *rrn3.8* | - | *ade5*, *his7-2*, *leu2-112*, *trp1-289*, *ura3-52*, *rrn3.8, GAL::HA-npa2 (kanMX6), RSA3::FPZ (NAT)* |
| *rrn3.8/RSA3::FPZ/ snr190-[mut.C]* | *rrn3.8* | - | *rrn3.8, snr190-[mut.C]* (CRISPR-Cas9), *RSA3::FPZ (NAT)* |
| *rrn3.8/RSA3::FPZ/ snr190-[mut.C]* | *rrn3.8* | pCH32 | *rrn3.8, snr190-[mut.C]* (CRISPR-Cas9), *RSA3::FPZ (NAT)* |
| *rrn3.8/RSA3::FPZ/ snr190-[mut.C]* | *rrn3.8* | pCH32 vector + wild-type *U14-SNR190* | *rrn3.8, snr190-[mut.C]* (CRISPR-Cas9), *RSA3::FPZ (NAT)* |
| *rrn3.8/RSA3::FPZ/ snr190-[mut.C]* | *rrn3.8* | pCH32 vector + *U14-snr190-[mut.AB]* | *rrn3.8, snr190-[mut.C]* (CRISPR-Cas9), *RSA3::FPZ (NAT)* |
| *rrn3.8/RSA3::FPZ/ snr190-[mut.C]* | *rrn3.8* | pCH32 vector + *U14-snr190-[mut.B]* | *rrn3.8, snr190-[mut.C]* (CRISPR-Cas9), *RSA3::FPZ (NAT)* |
| *rrn3.8/RSA3::FPZ/ snr190-[mut.C]* | *rrn3.8* | pCH32 vector + *U14-snr190-[short Δstem]* | *rrn3.8, snr190-[mut.C]* (CRISPR-Cas9), *RSA3::FPZ (NAT)* |
| *rrn3.8/RSA3::FPZ/ snr190-[mut.C]* | *rrn3.8* | pCH32 vector + *U14-snr190-[intermediate Δstem]* | *rrn3.8, snr190-[mut.C]* (CRISPR-Cas9), *RSA3::FPZ (NAT)* |
| *rrn3.8/RSA3::FPZ/ snr190-[mut.C]* | *rrn3.8* | pCH32 vector + *U14-snr190-[large Δstem]* | *rrn3.8, snr190-[mut.C]* (CRISPR-Cas9), *RSA3::FPZ (NAT)* |
| *rrn3.8/RSA3::FPZ/ snr190-[mut.C]* | *rrn3.8* | pCH32 vector + *U14-snr190-[mut.B-large Δstem]* | *rrn3.8, snr190-[mut.C]* (CRISPR-Cas9), *RSA3::FPZ (NAT)* |
| *NOP7::TAP/ snr190-[mut.C]* | BY4741 | pCH32 | *MATa*, *his3Δ1*, *leu2Δ0*, *met15Δ0*, *ura3Δ0*, *snr190-[mut.C]* (CRISPR-Cas9), *NOP7::TAP (HIS3)* |
| *NOP7::TAP/ snr190-[mut.C]* | BY4741 | pCH32 vector + wild-type *U14-SNR190* | *MATa*, *his3Δ1*, *leu2Δ0*, *met15Δ0*, *ura3Δ0*, *snr190-[mut.C]* (CRISPR-Cas9), *NOP7::TAP (HIS3)* |
| *NOP7::TAP/ snr190-[mut.C]* | BY4741 | pCH32 vector + *U14-snr190-[mut.AB]* | *MATa*, *his3Δ1*, *leu2Δ0*, *met15Δ0*, *ura3Δ0*, *snr190-[mut.C]* (CRISPR-Cas9), *NOP7::TAP (HIS3)* |
| *NOP7::TAP/ snr190-[mut.C]* | BY4741 | pCH32 vector + *U14-snr190-[mut.B]* | *MATa*, *his3Δ1*, *leu2Δ0*, *met15Δ0*, *ura3Δ0*, *snr190-[mut.C]* (CRISPR-Cas9), *NOP7::TAP (HIS3)* |
| *NOP7::TAP/ snr190-[mut.C]* | BY4741 | pCH32 vector + *U14-snr190-[short Δstem]* | *MATa*, *his3Δ1*, *leu2Δ0*, *met15Δ0*, *ura3Δ0*, *snr190-[mut.C]* (CRISPR-Cas9), *NOP7::TAP (HIS3)* |
| *NOP7::TAP/ snr190-[mut.C]* | BY4741 | pCH32 vector + *U14-snr190-[intermediate Δstem]* | *MATa*, *his3Δ1*, *leu2Δ0*, *met15Δ0*, *ura3Δ0*, *snr190-[mut.C]* (CRISPR-Cas9), *NOP7::TAP (HIS3)* |
| *NOP7::TAP/ snr190-[mut.C]* | BY4741 | pCH32 vector + *U14-snr190-[large Δstem]* | *MATa*, *his3Δ1*, *leu2Δ0*, *met15Δ0*, *ura3Δ0*, *snr190-[mut.C]* (CRISPR-Cas9), *NOP7::TAP (HIS3)* |
| *NOP7::TAP/ snr190-[mut.C]* | BY4741 | pCH32 vector + *U14-snr190-[mut.B-large Δstem]* | *MATa*, *his3Δ1*, *leu2Δ0*, *met15Δ0*, *ura3Δ0*, *snr190-[mut.C]* (CRISPR-Cas9), *NOP7::TAP (HIS3)* |

| | | | |
|---|---|---|---|
| W303 | - | - | *MAT*a, *leu2-3,112, trp1-1, can1-100, ura3-1, ade2-1, his3-11,15,* [phi+] |
| *snr190-[mut.C]* | W303 | - | *MAT*a, *leu2-3,112, trp1-1, can1-100, ura3-1, ade2-1, his3-11,15,* [phi+], *snr190-[mut.C]* (CRISPR-Cas9) |
| *snr190-[mut.C]/NOP8::FPZ* | W303 | - | *MAT*a, *leu2-3,112, trp1-1, can1-100, ura3-1, ade2-1, his3-11,15,* [phi+], *snr190-[mut.C]* (CRISPR-Cas9), *NOP8::FPZ (NAT)* |
| YAM1357 | W303 | pHT4467Δ-*NOP8* | *MAT*a, *leu2-3,112, trp1-1, can1-100, ura3-1, ade2-1, ade3*::*kanMX4*, *his3-11,15, nop8*::*HIS3MX4*, [phi+] |
| *NOP8::HTP* | YAM1357 | YCplac22-*NOP8*-HTP | *MAT*a, *leu2-3,112, trp1-1, can1-100, ura3-1, ade2-1, ade3*::*kanMX4*, *his3-11,15, nop8*::*HIS3MX4*, [phi+] |
| *Nop8ΔRRM-HTP* | YAM1357 | YCplac22-*NOP8.81C*-HTP | *MAT*a, *leu2-3,112, trp1-1, can1-100, ura3-1, ade2-1, ade3*::*kanMX4*, *his3-11,15, nop8*::*HIS3MX4*, [phi+] |
| *snr190-[mut.C]* | W303 | pCH32 | *MAT*a, *leu2-3,112, trp1-1, can1-100, ura3-1, ade2-1, his3-11,15,* [phi+], *snr190-[mut.C]* (CRISPR-Cas9) |
| *snr190-[mut.C]* | W303 | pCH32 vector + wild-type *U14-SNR190* | *MAT*a, *leu2-3,112, trp1-1, can1-100, ura3-1, ade2-1, his3-11,15,* [phi+], *snr190-[mut.C]* (CRISPR-Cas9) |
| *snr190-[mut.C]* | W303 | pCH32 vector + *U14-snr190-[mut.AB]* | *MAT*a, *leu2-3,112, trp1-1, can1-100, ura3-1, ade2-1, his3-11,15,* [phi+], *snr190-[mut.C]* (CRISPR-Cas9) |
| *snr190-[mut.C]* | W303 | pCH32 vector + *U14-snr190-[mut.B]* | *MAT*a, *leu2-3,112, trp1-1, can1-100, ura3-1, ade2-1, his3-11,15,* [phi+], *snr190-[mut.C]* (CRISPR-Cas9) |
| *snr190-[mut.C]* | W303 | pCH32 vector + *U14-snr190-[short Δstem]* | *MAT*a, *leu2-3,112, trp1-1, can1-100, ura3-1, ade2-1, his3-11,15,* [phi+], *snr190-[mut.C]* (CRISPR-Cas9) |
| *snr190-[mut.C]* | W303 | pCH32 vector + *U14-snr190-[intermediate Δstem]* | *MAT*a, *leu2-3,112, trp1-1, can1-100, ura3-1, ade2-1, his3-11,15,* [phi+], *snr190-[mut.C]* (CRISPR-Cas9) |
| *snr190-[mut.C]* | W303 | pCH32 vector + *U14-snr190-[large Δstem]* | *MAT*a, *leu2-3,112, trp1-1, can1-100, ura3-1, ade2-1, his3-11,15,* [phi+], *snr190-[mut.C]* (CRISPR-Cas9) |
| *snr190-[mut.C]* | W303 | pCH32 vector + *U14-snr190-[mut.B-large Δstem]* | *MAT*a, *leu2-3,112, trp1-1, can1-100, ura3-1, ade2-1, his3-11,15,* [phi+], *snr190-[mut.C]* (CRISPR-Cas9) |
| *GAL::HA-dbp6/ NOC1::FPZ* | MW3628 | - | *MATα, ura3-52, his3-Δ200, trp1-Δ63, leu2-Δ1, GAL::HA-dbp6 (kanMX6), NOC1::FPZ (NAT)* |
| *Δsnr37/snr190-[mutC]* | | pCH32 | *Δsnr37(NAT), snr190-[mut.C]* (CRISPR-Cas9) |
| *Δsnr37/snr190-[mutC]* | | pCH32 vector + wild-type *U14-SNR190* | *Δsnr37(NAT), snr190-[mut.C]* (CRISPR-Cas9) |
| *Δsnr37/snr190-[mutC]* | | pCH32 vector + *U14-snr190-[mut.AB]* | *Δsnr37(NAT), snr190-[mut.C]* (CRISPR-Cas9) |
| *Δsnr37/snR190-[mutC]* | | pCH32 vector + *U14-snr190-[mut.B]* | *Δsnr37(NAT), snr190-[mut.C]* (CRISPR-Cas9) |
| *Δsnr37/snR190-[mutC]* | | pCH32 vector + *U14-snr190-[short Δstem]* | *Δsnr37(NAT), snr190-[mut.C]* (CRISPR-Cas9) |
| *Δsnr37/snr190-[mutC]* | | pCH32 vector + *U14-snr190-[intermediate Δstem]* | *Δsnr37(NAT), snr190-[mut.C]* (CRISPR-Cas9) |
| *Δsnr37/snr190-[mutC]* | | pCH32 vector + *U14-snr190-[large Δstem]* | *Δsnr37(NAT), snr190-[mut.C]* (CRISPR-Cas9) |
| *Δsnr37/snr190-[mutC]* | | pCH32 vector + *U14-snr190-[mut.B-large Δstem]* | *Δsnr37(NAT), snr190-[mut.C]* (CRISPR-Cas9) |
